# Dual Immunoliposome Targeting of PD-L1 and CSF1R affects T-Cell readouts in tumor-conditioned co-cultures: An *In Vitro* Study in Glioblastoma and Medulloblastoma

**DOI:** 10.64898/2026.09.16.751417

**Authors:** Ommolbanin Asad Pour, Aylar Asadpour, Jamal Ghanam, Jan Best, Fatemeh Rahbarizadeh, Susann Hetze, Venkatesh Kumar Chetty, Lennart Barthel, Zohreh Amoozgar, Hartmut H Schmidt, Basant Kumar Thakur

## Abstract

Glioblastoma and medulloblastoma are characterized by an immunosuppressive tumor microenvironment, in which tumor-associated macrophages may impair T-cell function, in part through the expression of PD-L1 and CSF1R. Single-target therapies have shown inconsistent results in brain tumors, likely due to the complex interplay involving tumor-associated macrophages and immunosuppression. We developed two immunoliposomes functionalized with antibodies against PD-L1 and CSF1R to co-target these receptors on M2-like tumor-associated macrophages. TAM2Ms were generated by polarizing THP-1 monocytes with glioblastoma (A172, U87MG) and medulloblastoma (DAOY, ONS-76) tumor-conditioned medium together with IL-4 and IL-13. In co-culture with activated Jurkat T cells under tumor-conditioned medium-influenced conditions, combined αPD-L1 and αCSF1R immunoliposomes enhanced T-cell proliferation, reduced apoptosis, and improved migration toward tumor spheroids compared to single-target liposomal treatments or free antibodies. These benefits varied between glioblastoma and medulloblastoma in vitro models, reflecting distinct TAM2M phenotypes and microenvironment contexts. Dual immunoliposome targeting of PD-L1 and CSF1R may offer a promising strategy for reprogramming tumor-associated macrophages in brain tumor immunotherapy and warrants further evaluation in primary human macrophages and *in vivo* models to clarify efficacy and mechanisms.

**Graphical Abstract:** 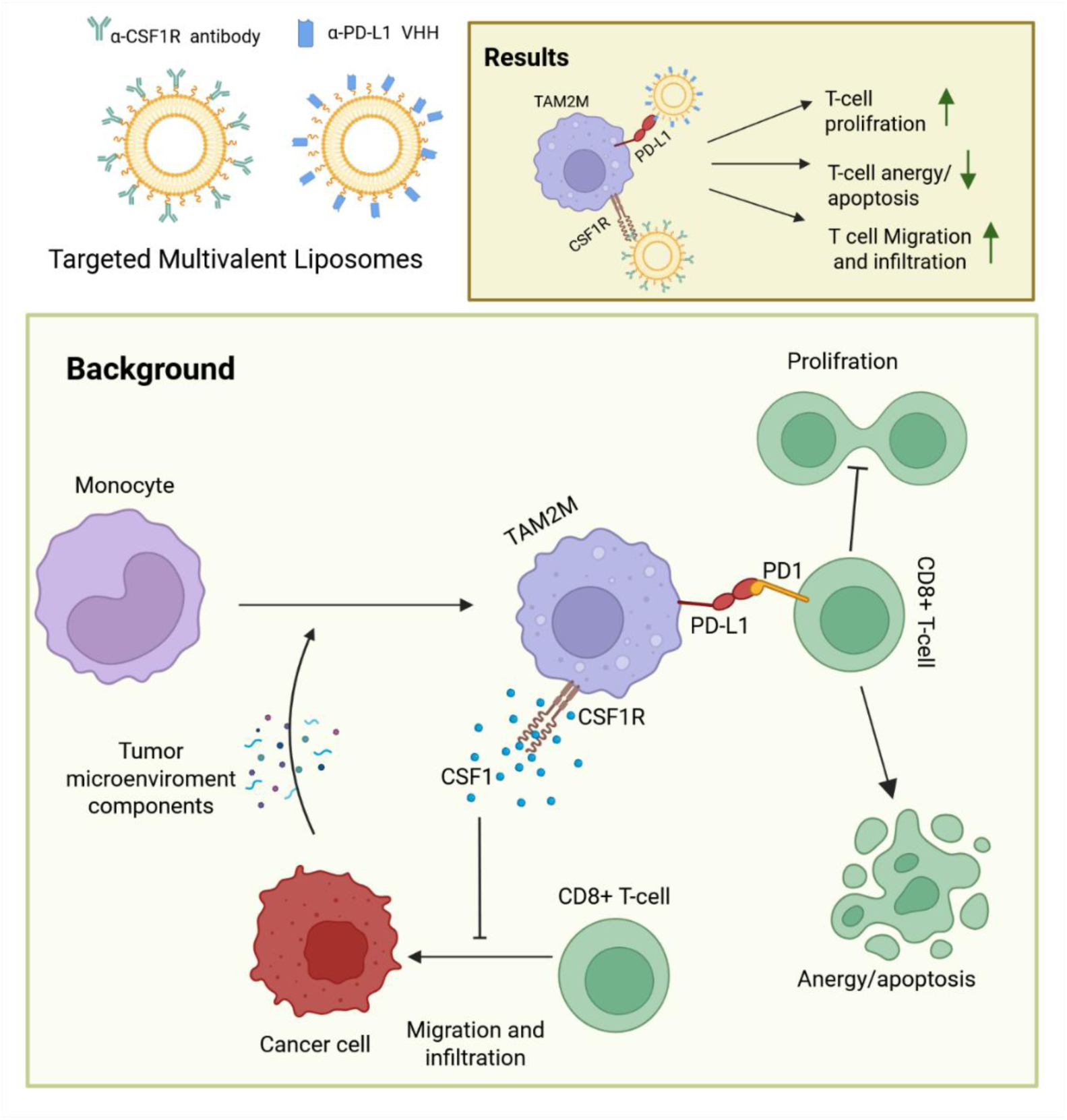

## Introduction

Brain tumors continue to pose a major challenge in oncology, contributing significantly to cancer-related illness and death [1]. Glioblastoma (GBM) is the most frequent malignant primary brain tumor in adults. It remains highly aggressive, with a median overall survival of only 14-16 months even with current standard-of-care treatments. These treatments typically involve surgical resection, followed by radiotherapy and chemotherapy with temozolomide [2–4]. Medulloblastoma (MB) is the most common malignant brain tumor in children and is associated with substantial treatment-related morbidity. Although survival has improved in certain molecular subgroups, recurrent or refractory disease remains a clinical challenge and is generally associated with poor prognosis [3,5,6]. The modest effectiveness of existing treatments for both GBM and MB highlights the urgent need for additional therapeutic approaches aimed at overcoming tumor recurrence and resistance to therapy.

The development of cancer immunotherapy, especially immune checkpoint blockade (ICB) targeting the programmed death-1/programmed death-ligand 1 (PD-1/PD-L1) axis, has resulted in meaningful improvements in outcomes for several solid tumor types [7], including melanoma [8], non-small cell lung cancer [9], and renal cell carcinoma [10] [11]. However, clinical trials evaluating PD-1/PD-L1 blockade in GBM and MB have yielded limited success, with low objective response rates and few patients experiencing durable benefits [12]. In recurrent GBM, the phase III Checkmate 143 trial failed to show a survival advantage for nivolumab compared with bevacizumab [13]. Similarly, the KEYNOTE-028 study reported an objective response rate of approximately 8% among PD-L1-positive recurrent GBM treated with pembrolizumab, highlighting the modest efficacy of single-agent checkpoint inhibition in these tumors [13,14]. In MB, clinical and translational data similarly indicate limited effectiveness of PD-1 blockade, with low response rates observed in unselected patient populations, suggesting that single-agent immune checkpoint inhibition may not be sufficient to overcome the immunosuppressive tumor microenvironment in this pediatric brain tumor [15,16].

The limited effectiveness of checkpoint blockade in brain tumors is likely due to multiple factors, but the immunosuppressive TME is a key contributor [17]. This environment can inhibit anti-tumor T-cell function and limit immunotherapy’s ability to elicit robust anti-tumor responses, thereby reducing treatment efficacy [18–20]. These findings have driven interest in combination strategies that pair checkpoint inhibitors with approaches to reprogram the tumor microenvironment and enhance T-cell activity, aiming to overcome immunosuppression and improve therapeutic outcomes in brain tumors [3,21].

A prominent characteristic of the tumor microenvironment is the substantial presence of tumor-associated macrophages (TAMs) [22,23]. In GBM, TAMs can account for 30–50% of the total tumor cellularity, making them a dominant immune population within the tumor and a key component of the immunosuppressive landscape [24–26]. Rather than promoting anti-tumor immunity, TAMs in brain tumors typically adopt an immunosuppressive M2-like phenotype, collectively known as tumor-associated M2 macrophages (TAM2Ms) [27,28]. Tumor-derived factors such as IL-10, TGF-β, CSF-1, IL-4, and IL-13 drive this polarization, shaping the microenvironment to support tumor progression and evade immune surveillance [29,30]. TAM2Ms suppress anti-tumor T-cell immunity through multiple interconnected mechanisms. These include the surface expression of immune checkpoint ligands such as PD-L1, which bind to inhibitory receptors on T cells and dampen their activity [31–34]. Programmed death-ligand 1 (PD-L1, CD274) is a transmembrane immune checkpoint molecule that is constitutively expressed on TAM2Ms and can be further induced by tumor-derived signals. When PD-L1 binds its receptor, PD-1, on T cells, it transmits co-inhibitory signals that dampen T-cell activation and proliferation and contribute to T-cell exhaustion and apoptosis. This interaction plays a key role in suppressing anti-tumor immune responses within the brain tumor microenvironment [32–34]. Although anti-PD-L1 antibody blockade has demonstrated potential to mitigate TAM-mediated suppression of T cells in preclinical models, its clinical use in brain tumors is constrained by several challenges [35,36]. These include the persistence of PD-L1-independent immunosuppressive pathways within the brain tumor microenvironment [37]. These factors contribute to the modest clinical efficacy observed in trials, highlighting the need for combination approaches to enhance therapeutic impact [31,36,38,39]. Colony-stimulating factor 1 receptor (CSF1R, CD115) is a receptor tyrosine kinase expressed on macrophages and monocytes and plays a key role in regulating their survival, proliferation, differentiation, and polarization toward an immunosuppressive M2 phenotype [40]. Activation of CSF1R by its ligands, CSF-1 and IL-34-both of which are secreted by brain tumor cells—promotes the recruitment and maintenance of tumor-associated macrophages (TAMs), particularly TAM2Ms, within the tumor microenvironment [41,42]. This signaling axis contributes significantly to establishing and sustaining an immunosuppressive niche that supports tumor progression [43]. In preclinical models, pharmacological inhibition of CSF1R has been shown to reduce TAM numbers or induce their repolarization toward a more pro-inflammatory M1-like phenotype, leading to partial restoration of anti-tumor immune responses and improved control of tumor growth [43,44]These findings support CSF1R as a promising target for modulating the tumor microenvironment in brain tumors [23,45]. Despite promising preclinical results, clinical trials of CSF1R inhibitors as monotherapy in glioblastoma (GBM) have yielded only modest clinical benefits [46]. This limited efficacy may be attributed to the activation of compensatory immunosuppressive mechanisms that operate independently of CSF1R signaling, thereby allowing the tumor microenvironment to maintain immune evasion despite macrophage depletion or repolarization [37]. These findings underscore the complexity of immune regulation in brain tumors and highlight the need for combination strategies that target multiple immunosuppressive pathways simultaneously [46].

These findings suggest that dual blockade of PD-L1 and CSF1R on tumor-associated M2 macrophages (TAM2Ms) may be more effective than targeting either pathway alone, as it simultaneously disrupts multiple immunosuppressive mechanisms driven by TAM2Ms [47,48]. This combination therapy via nanoparticle platforms-such as PEGylated immunoliposomes-offers several advantages over conventional systemic administration of free antibodies. These include multivalent antibody presentations on the nanoparticle surface. [49] for more effectively reprogramming the immunosuppressive tumor microenvironment in brain tumors [50,51]. Liposomal nanoparticles are a well-established and extensively studied drug delivery platform, with multiple formulations already approved for clinical use. [52–54] These systems have demonstrated favorable safety profiles and therapeutic efficacy in both preclinical models and human trials. [55]. PEGylated liposomes functionalized with targeting antibodies through thiol-maleimide receptor targeting on specific cell types-such as tumor-associated M2 macrophages-promote enhanced binding avidity, improved cellular uptake, and more effective modulation of target cell function. [54,56–58].

In this study, we developed and evaluated two antibody-functionalized PEGylated immunoliposomes, αPD-L1-Lipo and αCSF1R-Lipo, in *vitro*, designed to simultaneously target PD-L1 and CSF1R on tumor-associated M2 macrophages (TAM2Ms). These receptors were found to be co-expressed on TAM2Ms generated from THP-1 monocytes exposed to tumor-conditioned medium (TCM) derived from glioblastoma (GBM) cell lines (A172, U87MG) and medulloblastoma (MB) cell lines (DAOY, ONS-76). Using a series of co-culture assays with activated Jurkat T-cells and tumor spheroids, we showed that α PD-L1 and αCSF1R liposomes together significantly enhanced T-cell proliferation, reduced apoptosis, and improved T-cell migration toward tumor spheroids compared with single-target liposomal treatments or free antibodies. These effects were observed in a tumor microenvironment-dependent manner, highlighting the context-specific nature of TAM2M-mediated immunosuppression. Our findings provide *in vitro* evidence that dual immunoliposome targeting of TAM2Ms is a promising, rational strategy to overcome immunosuppression in both GBM and MB and warrants further investigation in more complex in vivo models [59,60].

## Methods

### Synthesis and characterization of αPD-L1-Lipo and αCSF1R-Lipo

#### Liposome Preparation

Antibody-functionalized liposomes (Fig. 1A) were prepared by lipid film hydration followed by extrusion. Briefly, 1,2-dioleoyl-sn-glycero-3-phosphoethanolamine (DOPE; 4.2 mg, 6 mmol; Avanti Polar Lipids, AL, USA ), cholesteryl hemisuccinate (CHEMS; 1.94 mg, 4 mM; Sigma-Aldrich, St. Louis, USA), 1,2-distearoyl-sn-glycero-3-phosphoethanolamine-N-[methoxy (polyethylene glycol)-2000] (mPEG2000-DSPE; 0.17 mg, 0.06 mM; ChemScene, NJ, USA), and 1,2-distearoyl-sn-glycero-3-phosphoethanolamine-N-[maleimide (polyethylene glycol)-2000] (DSPE-PEG (2000)-Maleimide; 0.48 mg, 0.24 mM; MedChemExpress, NJ, USA ) were dissolved in 1 mL of chloroform at the molar ratios indicated above. The organic solvent was evaporated under reduced pressure using a rotary evaporator to form a thin, uniform lipid film. The dried film was then hydrated with phosphate-buffered saline (PBS, pH 7.4) to generate self-assembled multilamellar vesicles. The resulting liposome suspension was probe-sonicated using an ultrasonic homogenizer (UP400s; Hielscher Ultrasonics, Teltow, Germany) at 130 W and 40 kHz for 60 seconds (two 30-second cycles with 10-second intervals) at 25°C to reduce vesicle size. The suspension was subsequently extruded 11 times through 0.1 µm polycarbonate membranes (Avanti Research, Al, USA) using a hand-held mini-extruder (Avanti Research, Al, USA) to produce unilamellar PEGylated liposomes (PEG-Lipo) with a defined size distribution. PEG-Lipo served as the base formulation for subsequent antibody conjugation.

**Figure 1.**
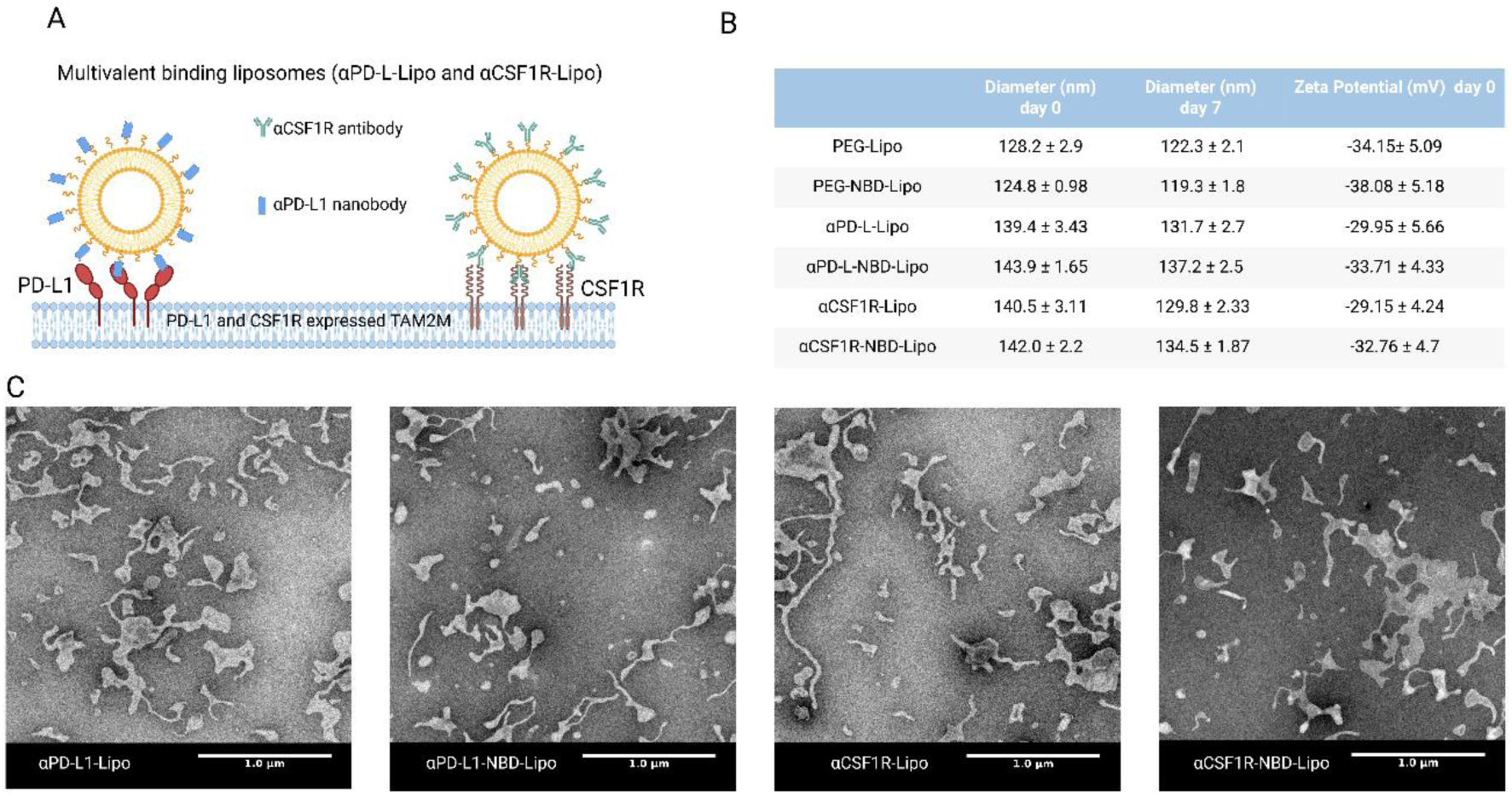
Design, physicochemical characterization, and morphological assessment of antibody-functionalized multivalent binding liposomes. (A) Schematic illustration of the two targeted liposomal formulations: αPD-L1-Lipo, conjugated with anti-PD-L1 nanobody (αPD-L1 Nb, blue rectangles) and αCSF1R-Lipo functionalized with anti-CSF1R antibody (Cabiralizumab, green Y-shaped antibodies), designed to engage PD-L1 and CSF1R receptors, respectively, on the surface of tumor-associated M2 macrophages (TAM2Ms). (B) Physicochemical characterization table summarizing mean hydrodynamic diameter (nm) at day 0 and day 7 of storage at 4°C, and zeta potential (mV) at day 0, for all liposomal formulations: PEG-Lipo, PEG-NBD-Lipo, αPD-L1-Lipo, αPD-L1-NBD-Lipo, αCSF1R-Lipo, and αCSF1R-NBD-Lipo. Data are expressed as mean ± SD (n = 3). (C) Representative transmission electron microscopy (TEM) micrographs of negatively stained αPD-L1-Lipo, αPD-L1-NBD-Lipo, αCSF1R-Lipo, and αCSF1R-NBD-Lipo formulations. Scale bar = 1.0 µm.

#### Antibody Thiolation and Conjugation

To functionalize liposomes with targeting antibodies, anti-PD-L1 nanobody (αPD-L1 Nb; provided by the Opazo lab (https://opazolab.de/) and anti-CSF1R antibody (Cabiralizumab; synonyms: FPA 008; anti-Human CSF1R recombinant antibody; MedChemExpress, NJ, USA) were each thiolated using Traut’s reagent (2-iminothiolane hydrochloride; Sigma-Aldrich, St. Louis, MO, USA) at a concentration of 2 mg/mL for 60 minutes at 27°C with gentle agitation, to introduce free thiol groups onto the antibody surface. The thiolated antibodies were then conjugated to maleimide-functionalized PEG-Lipo via thiol-maleimide chemistry, forming stable thioether linkages between the antibodies and the liposome surface. Excess unreacted antibody was removed by size-exclusion chromatography using a Sepharose CL-4B column equilibrated with PBS. For fluorescence-based tracking of liposomal uptake *in vitro*, NBD-PE fluorescent lipid was incorporated into the lipid mixture at a concentration of 0.06 mM (1 mol% of the total phosphatidylethanolamine content) during film preparation, yielding fluorescently labeled NBD-αPD-L1-Lipo and NBD-αCSF1R-Lipo formulations for cellular uptake studies.

#### Physicochemical Characterization

The hydrodynamic size distribution and surface charge of all liposomal formulations were determined by nanoparticle tracking analysis (NTA) using a ZetaView BASIC PMX-120 instrument (Particle Metrix GmbH, Inning am Ammersee, Germany). We assessed liposome availability by transmission electron microscopy (TEM; JEOL Ltd., Tokyo, Japan) after negative staining with phosphotungstic acid (PTA). We quantified the concentration of conjugated αPD-L1 Nb and αCSF1R Ab in purified liposomal preparations using the bicinchoninic acid (BCA) protein assay according to the manufacturer’s instructions. Antibody conjugation efficiency was calculated using the following formula:

Conjugation efficiency (%) = (amount of antibody conjugated to the liposomes/initial amount of antibody added) *100

To confirm successful surface conjugation of αPD-L1 Nb and αCSF1R Ab to the liposomes, sodium dodecyl sulfate-polyacrylamide gel electrophoresis (SDS-PAGE) was performed under reducing conditions on a 4-20% polyacrylamide gel, and the proteins were subsequently visualized by Coomassie Brilliant Blue R-250 staining.

#### Stability Assessment of αPD-L1-Lipo and αCSF1R-Lipo

The colloidal stability of αPD-L1-Lipo and αCSF1R-Lipo formulations was evaluated by monitoring changes in mean hydrodynamic particle size over time. Formulations were stored at 4°C, and particle sizes were measured by NTA at baseline (day 0) and following 7 days of storage.

### Preparation of Tumor-Conditioned Medium (TCM)

To generate TCM, glioblastoma (GBM) cell lines (U-87MG and A-172) and medulloblastoma (MB) cell lines (ONS-76 and DAOY) were each cultured to approximately 80% confluence in their respective standard growth media. At this point, the culture medium was replaced with RPMI-1640 supplemented with 1% FBS, and cells were incubated for 48 hours at 37°C in 5% CO₂. Following the conditioning period, the supernatant was collected and centrifuged at 200 × g for 10 min at 4°C to remove cellular debris and non-adherent cells. The clarified supernatant was then passed through a 0.45 µm syringe filter to ensure sterility, and FBS was added to a final concentration of 9% (v/v) to restore physiologically relevant serum conditions. We stored TCM aliquots at −80°C until use.

### Differentiation and polarization of THP-1 Monocytes into Tumor-Associated M2 Macrophages (TAM2Ms)

THP-1 human monocytic cells were differentiated and polarized into tumor-associated M2 macrophages (TAM2Ms) as previously described in our manuscript [61], with minor modifications. Briefly, THP-1 cells were seeded at a density of 3×10^5^ cells per well in 24-well plates in RPMI-1640 supplemented with 10% heat-inactivated FBS (HI-FBS) and treated with 150 ng/mL phorbol 12-myristate 13-acetate (PMA; Sigma-Aldrich) for 48 hours at 37°C in a humidified atmosphere containing 5% CO₂ to induce macrophage differentiation. After differentiation, remove the PMA-containing medium, wash adherent cells once with fresh RPMI-1640 supplemented with 10% HI-FBS, and allow them to rest for 24 hours. To induce TAM2M polarization, differentiated macrophages were subsequently cultured for 72 hours in a 1:1 (v/v) mixture of RPMI-1640 supplemented with 10% HI-FBS and tumor-conditioned medium (TCM) derived from each respective tumor cell line, further supplemented with recombinant human IL-4 and IL-13 (25 ng/mL each; PeproTech) (Fig. S3A). Morphological changes associated with macrophage differentiation and polarization were documented by phase-contrast imaging using an inverted fluorescence microscope (EVOS FL; Thermo Fisher Scientific, MA, USA) at 10× magnification under standardized light intensity settings. We acquired representative images from randomly selected fields across the entire well area.

### Cytotoxicity Assessment of Targeted Liposomes on TAM2Ms

The cytotoxic potential of PEG-Lipo, αPD-L1-Lipo, and αCSF1R-Lipo on TAM2M-polarized THP-1 macrophages was evaluated using the Cell Counting Kit-8 (CCK-8; MedChemExpress, NJ, USA) colorimetric assay. We seeded THP-1 cells at 3 × 10⁴ cells per well in 96-well plates, then differentiated and polarized them into TAM2Ms as described above. After polarization, cells were pretreated with 1 µM cytochalasin D (Millipore Sigma, Burlington, MA, USA) for 2 hours at 37°C to inhibit phagocytic uptake of liposomes and ensure that the observed effects reflected receptor-mediated interactions rather than nonspecific engulfment. Cells were subsequently treated with serial dilutions of each liposomal formulation (0–10 mg/mL of conjugated antibody equivalent) for 48 hours at 37°C in 5% CO₂. Following the treatment period, the culture medium was replaced with fresh medium containing 10% (v/v) CCK-8 solution, and cells were incubated for 60 minutes at 37°C. Absorbance was measured at 450 nm using a microplate reader (Tecan Nano Quant Infinite M200 Pro; Tecan Group Ltd., Männedorf, Switzerland). Cell viability was expressed as a percentage relative to untreated control cells, and data were analyzed using GraphPad Prism (version 11; GraphPad Software, San Diego, CA, USA).

### *In Vitro* Cellular Internalization Assay

To evaluate receptor-mediated internalization of αPD-L1-Lipo and αCSF1R-Lipo by TAM2Ms, THP-1 cells were seeded at 3 × 10⁵ cells per well in 24-well plates, then differentiated and polarized into TAM2Ms as described above. Following polarization, cells were pre-treated with 1 µM cytochalasin D for 2 hours. Cells were then incubated with fluorescently labeled NBD-αPD-L1-Lipo or NBD-αCSF1R-Lipo for 24 hours at 37°C in 5% CO₂. Non-targeted-PEGylated liposomes lacking conjugated antibodies (NBD-PEG-Lipo) served as a control to distinguish receptor-mediated uptake from passive association. Following incubation, cells were washed twice with PBS to remove unbound liposomes, fixed with 4% paraformaldehyde for 15 minutes at room temperature, and counterstained with DAPI (300 nM) to visualize nuclei. We acquired fluorescence images at 20× magnification using an inverted fluorescence microscope and quantified intracellular fluorescence intensity using ImageJ software (version 1.54; National Institutes of Health, Bethesda, MD, USA).

### Flow Cytometric Analysis of PD-L1 and CSF1R Surface Expression on TAM2Ms

Surface expression of PD-L1 (CD274) and CSF1R (CD115) on TAM2M-polarized THP-1 cells was quantified by flow cytometry. We seeded THP-1 cells at 3 × 10⁵ cells per well in 24-well plates and polarized them into TAM2Ms as described above. Cells were harvested using Versene (0.5 mM EDTA in PBS; 8 minutes at 37°C), washed twice with ice-cold staining buffer (PBS supplemented with 2% BSA and 0.02% sodium azide), and counted. To block non-specific antibody binding, incubate cells with 5 µL of Human TruStain FcX™ Fc Receptor Blocking Solution (BioLegend, San Diego, CA, USA) per 100 µL cell suspension for 10 minutes at room temperature. Cells were then incubated on ice for 20 minutes with primary antibodies against CSF1R (anti-human CSF1R recombinant antibody; MedChemExpress, NJ, USA) and PD-L1 (anti-human CD274/PD-L1/B7-H1, clone MIH1; eBioscience™, Thermo Fisher Scientific, MA, USA), each at a 1:1000 dilution in staining buffer. Following primary antibody incubation, cells were washed twice with staining buffer by centrifugation at 350 × g for 5 minutes, then incubated with species-appropriate FITC-conjugated secondary antibodies (anti-rabbit and anti-mouse IgG; 1:5,000 dilution) for 15 minutes on ice in the dark. Fluorescence data were acquired using a CytoFLEX flow cytometer (Beckman Colter, Suzhou, China), and mean fluorescence intensity (MFI) values were quantified using the online Flow Cytometry Analysis Tool (https://floreada.io/). Statistical analyses were performed using GraphPad Prism (version 11.6.1; GraphPad Software, San Diego, CA, USA).

### Flow cytometric analysis of M2 macrophage marker expression

To confirm the M2 polarization phenotype of TAM2Ms, we seeded THP-1 cells at 3 × 10⁵ cells per well in 24-well plates and polarized them into TAM2Ms as described above. Following polarization, cells were washed with PBS and harvested using Versene. Cell pellets were resuspended in staining buffer (PBS supplemented with 1% FBS), and Fc receptors were blocked using 5 µL of Human TruStain FcX™ per 89 µL cell suspension for 10 minutes at room temperature. Cells were then stained with anti-CD163-APC (2 µL) for 20 minutes in the dark at 2–8°C, washed twice with staining buffer, and resuspended for acquisition. Data were acquired using a CytoFLEX flow cytometer, and analysis was performed using the online Flow Cytometry Analysis Tool. Statistical analyses were carried out using GraphPad Prism.

### Assessment of T-Cell Proliferation by CFSE Dilution Assay

We assessed the effect of pretreated TAM2Ms on Jurkat T-cell proliferation using a CFSE dilution assay monitored by flow cytometry. Jurkat T-cells were labeled with CellTrace™ CFSE (Invitrogen, Thermo Fisher Scientific, MA, USA) by dissolving the reagent in DMSO and diluting it in pre-warmed PBS (37°C). To quench excess dye, add 40 mL of complete RPMI-1640 and incubate for 10 minutes at 37°C with gentle inversion every 2 minutes. Cells were collected by centrifugation at 300 × g for 5 minutes. For polyclonal activation, Dynabeads™ Human T-Activator CD3/CD28 (Thermo Fisher Scientific) were used at a bead-to-cell ratio of 1:3 (6.7 × 10⁶ beads per 2 × 10⁷ cells). Beads were washed by magnetic separation, resuspended in complete RPMI-1640 supplemented with 10% FBS and recombinant human IL-2 (10 ng/mL), and added to the CFSE-labeled T-cell suspension. Cells were incubated for 24 hours at 37°C in 5% CO₂. Following activation, beads were removed by magnetic separation, and activated Jurkat T-cells were washed twice with RPMI-1640 supplemented with 10% HI-FBS. Cells were then co-cultured with TAM2Ms pretreated with PEG-Lipo, αPD-L1, αPD-L1-Lipo, αCSF1R, and αCSF1R-Lipo, or a combination of αPD-L1-Lipo and αCSF1R-Lipo at a 1:1 effector-to-target ratio for 72 hours at 37°C in 5% CO₂. T-cell proliferation was assessed by flow cytometry using sequential CFSE dilution, whereby each cell division results in a twofold reduction in fluorescence intensity. Data acquisition was performed on a CytoFLEX flow cytometer, and proliferation index calculation was performed using the online Flow Cytometry Analysis Tool.

### Flow Cytometric Analysis of T-Cell Apoptosis by Annexin V/Propidium Iodide Staining

T-cell apoptosis was assessed by dual staining with Annexin V-FITC and propidium iodide (PI) to distinguish viable, early apoptotic, late-apoptotic, and necrotic cell populations. We maintained Jurkat T-cells in RPMI-1640 supplemented with 10% FBS and recombinant human IL-2 (10 ng/mL) throughout the THP-1 differentiation and polarization period. Twenty-four hours before co-culture, Jurkat T-cells were polyclonally activated using Dynabeads™ Human T-Activator CD3/CD28 as described above. Following activation and bead removal, cells were resuspended in complete medium and co-cultured with TAM2Ms pretreated with PEG-Lipo, αPD-L1, αPD-L1-Lipo, αCSF1R, and αCSF1R-Lipo, or a combination of αPD-L1-Lipo and αCSF1R-Lipo, for 72 hours at 37°C in 5% CO₂. Following co-culture, Jurkat T-cells were harvested by centrifugation at 300 × g for 5 minutes, washed once with ice-cold PBS, and stained using the Annexin V-FITC Apoptosis Detection Kit (ab14085; Abcam, Cambridge, UK) according to the manufacturer’s instructions. Briefly, cells were resuspended in binding buffer and incubated with Annexin V-FITC and PI for 15 minutes at room temperature in the dark.

Samples were acquired immediately using a CytoFLEX flow cytometer. Cell populations were distinguished as follows: viable cells (Annexin V⁻/PI⁻), early apoptotic cells (Annexin V⁺/PI⁻), late apoptotic/secondary necrotic cells (Annexin V⁺/PI⁺), and necrotic cells (Annexin V⁻/PI⁺). Data were analyzed using the online Flow Cytometry Analysis Tool and GraphPad Prism.

### Jurkat T-Cell Migration Toward Tumor Spheroids

We performed a transwell-based migration assay using three-dimensional GBM and MB spheroid models, as described below.

#### Tumor Spheroid Formation

Three-dimensional tumor spheroids were generated by seeding 2,500 GBM (U-87MG, A-172) or MB (ONS-76, DAOY) cells per well in Millicell® Ultra-low Attachment Plates (Millicell, Darmstadt, Germany) in 100 µL of serum-free spheroid medium. Spheroid medium consisted of DMEM supplemented with 2% B-27 Supplement (50x, minus Vitamin A; Gibco, New York, USA), 1× N2 supplement (Gibco, New York, USA), 2 mM L-glutamine, 20 ng/mL recombinant human EGF, 20 ng/mL recombinant human FGF-2, and 1% Penicillin/Streptomycin. Spheroids were allowed to self-assemble over 48 hours at 37°C in a humidified 5% CO₂ atmosphere and were confirmed and imaged by inverted microscopy before use in migration assays.

#### Jurkat T-Cell Fluorescent Labeling and Activation

Jurkat T-cells were fluorescently labeled with CellTrace™ CFSE and polyclonally activated as described above.

#### Co-culture of Activated T Cells with Pretreated TAM2Ms

CFSE-labeled, activated Jurkat T-cells were co-cultured with TAM2Ms pretreated with PEG-Lipo, αPD-L1, αPD-L1-Lipo, αCSF1R, and αCSF1R-Lipo or a combination of αPD-L1-Lipo and αCSF1R-Lipo at a 1:1 ratio for 72 hours at 37°C in 5% CO₂ before use in migration experiments.

**Transwell Migration Assay Setup: Cultrex Basement Membrane Extract (BME; 8–12 mg/mL; R&D Systems by Bio-Techne, Nordstadt,** Germany) was thawed overnight at 2–8°C to prevent premature gelation, as BME solidifies irreversibly above 15°C. All handling was performed on ice, and BME was gently homogenized by slow pipetting to avoid air bubbles. We carefully dispensed 120 µL of BME into each pre-chilled Millicell® 24-well standing insert (pore size 8.0 μm, diameter 12 mm, Millipore, Darmstadt, Germany) and spread it evenly across the membrane. Incubate coated inserts at 37°C for 30 minutes to allow complete matrix gelation. T cells were gently dislodged using cold PBS supplemented with 2–5 mM EDTA, washed once with RPMI-1640, resuspended at the appropriate density in 100 µL of complete medium, and loaded into the upper chamber of the BME-coated Transwell inserts. Place pre-formed tumor spheroids in the lower chamber to create a chemoattractant gradient that drives directional T-cell migration through the BME matrix.

#### Assessment of Jurkat T-Cell Migration

Following 48 hours of incubation at 37°C in 5% CO₂, Transwell inserts were carefully removed, and the membranes were gently washed with PBS to remove non-migrated cells and residual medium. CFSE-labeled T cells that had migrated through the BME matrix were visualized and enumerated by an epifluorescence microscope and quantified using ImageJ software. We estimated the total number of migrated CFSE⁺ T cells across the entire Transwell membrane surface area by calculating the total image area from pixel dimensions, dividing the total membrane area by the single-image area to determine the number of fields of view, and multiplying this value by the mean cell count per image.

## Results

### Synthesis and Physicochemical Characterization of Antibody-Functionalized Liposomal Nanoparticles

Nanoparticle size and surface charge are critical determinants of multivalent receptor targeting, as they influence cellular uptake and biodistribution. The non-functionalized PEGylated base formulation (PEG-Lipo) exhibited a mean hydrodynamic diameter of 128.2 ± 2.9 nm on day 0 and 122.3 ± 2.1 nm on day 7, indicating good colloidal stability with no evidence of particle growth or aggregation over the storage period. The fluorescent analog (PEG-NBD-Lipo) showed comparable size and stability (124.8 ± 0.98 nm on day 0; 119.3 ± 1.8 nm on day 7), demonstrating that incorporation of NBD-PE did not alter particle behavior.

Surface conjugation with targeting antibodies resulted in the expected modest increase in particle size. αPD-L1-Lipo and αPD-L1-NBD-Lipo measured 139.4 ± 3.43 nm and 143.9 ± 1.65 nm, respectively, on day 0 and remained stable over 7 days (131.7 ± 2.7 nm and 137.2 ± 2.5 nm). Similarly, αCSF1R-Lipo and αCSF1R-NBD-Lipo measured 140.5 ± 3.11 nm and 142.0 ± 2.2 nm on day 0 and 129.8 ± 2.33 nm and 134.5 ± 1.87 nm on day 7.

Zeta potentials on day 0 ranged from −29.15 ± 4.24 mV (αCSF1R-Lipo) to −38.08 ± 5.18 mV (PEG-NBD-Lipo), reflecting a consistently negative surface charge attributable to the anionic lipid components (CHEMS) and PEG coating. This surface charge profile is consistent with electrostatic stabilization and supports the observed colloidal stability across all formulations (Fig. 1B; Fig. S1). Number-based size distributions by NTA were unimodal across all formulations and shifted slightly toward smaller sizes by day 7, with zeta-potential histograms centered between approximately −30 and −40 mV (Fig. S1).

TEM imaging of αPD-L1-Lipo, αPD-L1-NBD-Lipo, αCSF1R-Lipo, and αCSF1R-NBD-Lipo revealed nanoscale vesicles distributed across the imaging field (scale bar, 1.0 µm). Particles appeared flattened/irregular on the grid, which is consistent with adsorption and partial dehydration of flexible phospholipid bilayers during TEM sample preparation and does not reflect the native solution-phase morphology of the liposomes (Fig. 1C).

Biochemical confirmation of successful surface conjugation was obtained by reducing SDS-PAGE (Fig. S2). No protein bands were detected in the PEG-Lipo lane, confirming the absence of protein in the non-functionalized formulation. The free αPD-L1 nanobody (αPD-L1 Nb) displayed a major band at approximately 17 kDa with a faint dimer band near ∼32 kDa; corresponding bands at the same positions were observed in the αPD-L1-Lipo lane, confirming successful conjugation of the nanobody to the liposomal surface. The free αCSF1R antibody (Cabiralizumab) exhibited the expected IgG heavy-chain (∼50 kDa) and light-chain (∼25 kDa) bands, which were also present in the αCSF1R-Lipo lane, verifying successful coupling of the antibody to the nanoparticle surface. We used a pre-stained molecular weight marker (10–150 kDa) and Coomassie Brilliant Blue staining for molecular weight reference and protein visualization, respectively.

### Tumor-Conditioned Medium Upregulates PD-L1, CSF1R, and CD163 Expression on THP-1-Derived Macrophages, Confirming TAM2M Polarization

To verify co-expression of the intended therapeutic targets on polarized macrophages, surface levels of PD-L1 and CSF1R were quantified by mean fluorescence intensity (MFI) on THP-1–derived macrophages cultured with each tumor-conditioned medium (TCM), and CD163 MFI was measured across the polarization hierarchy (Fig. 2A) Relative to unpolarized M0 controls, PD-L1 surface expression was significantly elevated under all four TCM conditions, with A172-TCM (p < 0.05), U87MG-TCM (p < 0.01), DAOY-TCM (p < 0.05) and ONS-76-TCM (p < 0.05) each producing higher MFI values. CSF1R expression showed a more differential pattern: A172-TCM produced a non-significant trend relative to M0 control, whereas U87MG-TCM, DAOY-TCM and ONS-76-TCM each resulted in significant upregulation (all p < 0.05).

**Figure 2.**
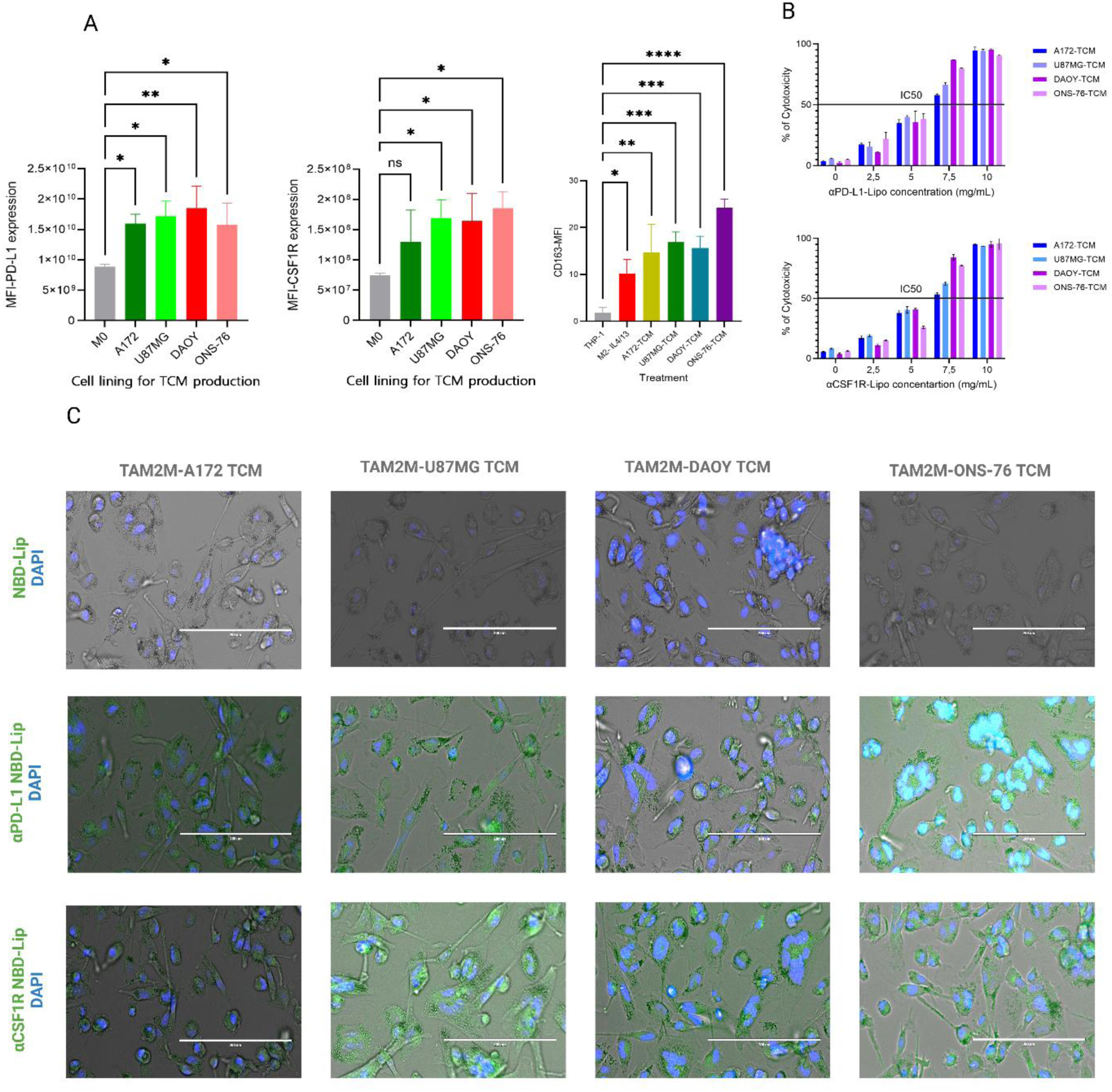
Tumor-conditioned medium drives upregulation of PD-L1, CSF1R, and CD163 on THP-1-derived macrophages, confirming successful TAM2M polarization. (A) Flow cytometric quantification of PD-L1 MFI (left) and CSF1R MFI (center) on THP-1-derived macrophages maintained under unpolarized conditions (M0) or polarized with TCM derived from A172, U87MG, DAOY, or ONS-76 tumor cell lines supplemented with IL-4 and IL-13 (25 ng/mL each). CD163 MFI (Right) across undifferentiated THP-1 monocytes, IL-4/IL-13-polarized macrophages (M2-IL4/13), and TAM2Ms polarized under A172-TCM, U87MG-TCM, DAOY-TCM, and ONS-76-TCM conditions. Data are expressed as mean ± SEM (n = 3). Statistical significance was determined by one-way ANOVA with Tukey’s post-hoc test (*p < 0.05, **p < 0.01, ***p < 0.001, ****p < 0.0001; ns = not significant). (B) Concentration-dependent cytotoxicity of αPD-L1-Lipo (top) and αCSF1R-Lipo (bottom) on TAM2Ms polarized under A172-TCM, U87MG-TCM, DAOY-TCM, and ONS-76-TCM conditions. The horizontal dashed line indicates the IC50 (50% cytotoxicity) threshold. Data are expressed as mean ± SEM (n = 3). (C) Representative fluorescence microscopy images of TAM2Ms incubated with NBD-Lipo (non-targeted control), αPD-L1-NBD-Lipo, or αCSF1R-NBD-Lipo. Green channel: NBD fluorescence (liposome internalization); Blue channel: DAPI nuclear stain. Images were acquired at 20× magnification. Scale bar 200µm.

CD163 expression increased progressively with advancing polarization state (Fig. 2A, right). Undifferentiated THP-1 monocytes exhibited the lowest CD163 MFI. Treatment with IL-4/IL-13 alone (M2-IL4/13) modestly but significantly elevated CD163 compared to monocytes (p < 0.05). The addition of TCM further enhanced CD163 expression beyond that achieved with IL-4/IL-13 alone, with A172-, U87MG- and DAOY-TCM each yielding significant increases (all p < 0.001), and ONS-76-TCM producing the highest CD163 MFI of all conditions (p < 0.0001 vs monocytes), collectively indicating robust TAM2M polarization. Gating strategies and representative flow cytometry plots supporting these marker analyses are provided in Figures S2–S3.

### Cytotoxicity of αPD-L1-Lipo and αCSF1R-Lipo on TAM2M

Dose–response CCK-8 assays demonstrated minimal cytotoxicity for both targeted liposomal formulations at concentrations between 0 and 2.5 mg/mL across all TCM-polarized TAM2M conditions, followed by a progressive increase in cytotoxicity between 2.5 and 7.5 mg/mL. The IC50 region was observed at approximately 5–7.5 mg/mL, depending on the TCM condition (Fig. 2B). At the highest tested concentration of 10 mg/mL, cytotoxicity exceeded 80–100% across all conditions. Importantly, the working dose used in downstream co-culture experiments (5 mg/mL) remained well below the IC50 for all TCM conditions and both liposomal formulations, supporting its suitability for functional assays.

### Receptor-Mediated Cellular Internalization of Targeted Liposomes by TAM2Ms

Fluorescence microscopy confirmed target-specific uptake of antibody-functionalized liposomes by TAM2Ms (Fig. 2C). Non-targeted NBD-PEG-Lipo produced little to no detectable intracellular fluorescence signal in TAM2Ms generated under any TCM condition, indicating negligible passive association. In contrast, both αPD-L1-NBD-Lipo and αCSF1R-NBD-Lipo produced strong punctate cytoplasmic fluorescence across all TCM conditions, consistent with active receptor-mediated internalization rather than nonspecific surface binding or passive uptake.

### TAM2M Pretreatment with Targeted Immunoliposomes Restores Jurkat T-cell Proliferation in a TCM-dependent Manner

CFSE-labeled, anti-CD3/CD28–activated Jurkat T-cells were co-cultured with TAM2Ms generated using TCM from A172, U87MG, DAOY, or ONS-76 cell lines. T-cell proliferation was quantified by CFSE dilution and compared across seven conditions: untreated, PEG-Lipo, free αPD-L1, free αCSF1R, αPD-L1-Lipo, αCSF1R-Lipo, and the combination αPD-L1+αCSF1R-Lipo (Fig. 3)

**Figure 3.**
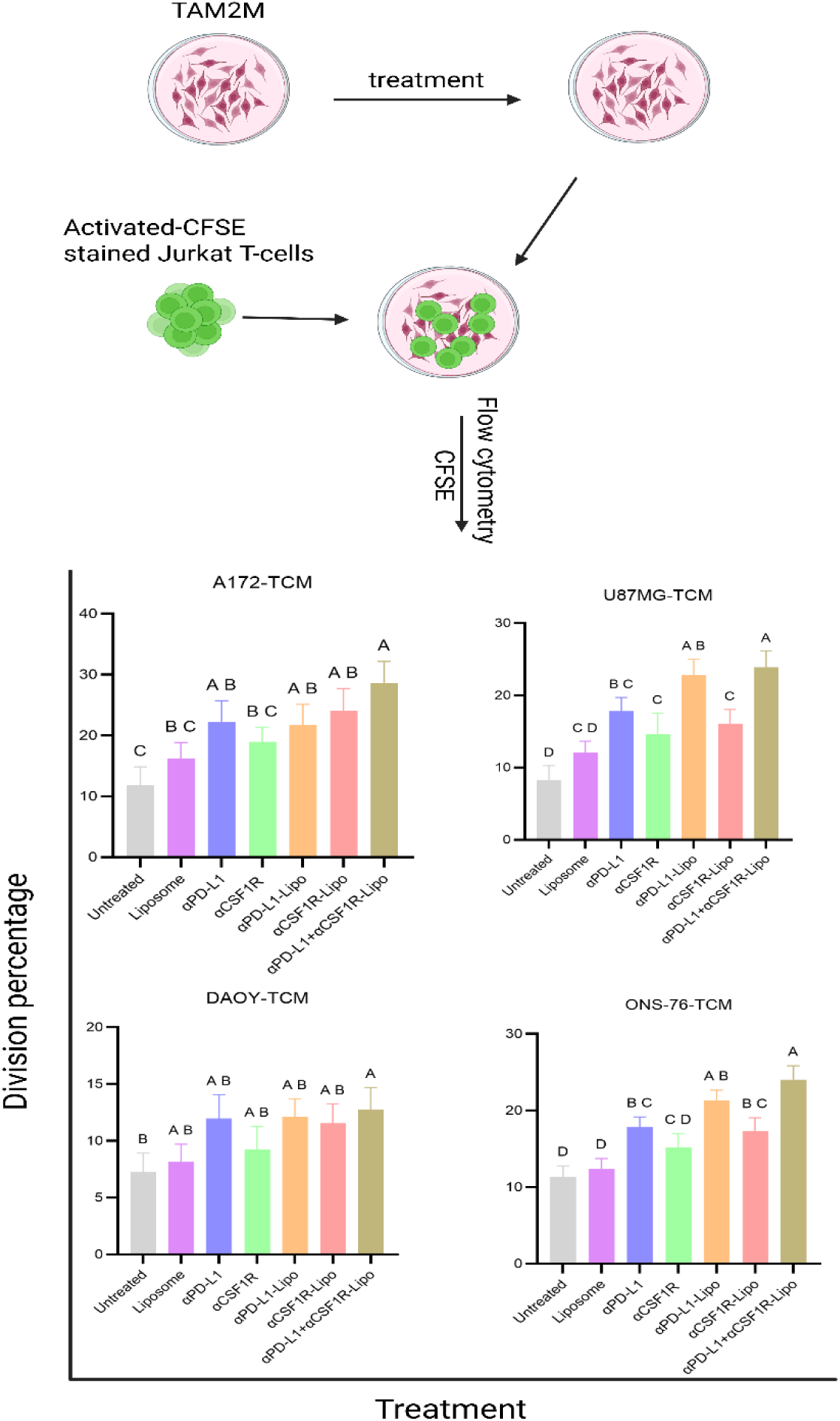
Liposomal co-targeting of PD-L1 and CSF1R on TAM2Ms restores Jurkat T-cell proliferation across GBM and MB tumor-conditioned medium models. (Top) Schematic representation of the experimental workflow. (Bottom) Bar graphs depicting the T-cell division percentage following co-culture with TAM2Ms pretreated under seven conditions: Untreated, PEG-Lipo (empty liposome control), free αPD-L1 nanobody (αPD-L1), free αCSF1R antibody (αCSF1R), αPD-L1-Lipo, αCSF1R-Lipo, and dual combination αPD-L1-Lipo+αCSF1R-Lipo. Results are shown for TAM2Ms generated under A172-TCM, U87MG-TCM, DAOY-TCM, and ONS-76-TCM conditions. Data are presented as mean ± SEM (n = 3). Different letters above bars (A, B, C, D) indicate statistically significant differences between groups as determined by one-way ANOVA with Tukey’s post-hoc test (p < 0.05). Groups sharing the same letter are not significantly different from one another.

#### A172-TCM TAM2Ms

Untreated co-cultures exhibited the lowest level of T-cell division. PEG-Lipo did not significantly alter proliferation relative to untreated controls. Free αPD-L1 increased T-cell division (AB vs C), whereas free αCSF1R showed only a nonsignificant trend toward improvement (BC vs C). Single-targeted liposomal formulations (αPD-L1-Lipo and αCSF1R-Lipo) both ranked in the top statistical group (AB). The dual αPD-L1+αCSF1R-Lipo combination yielded the highest mean division (A) and was significantly greater than untreated, PEG-Lipo, and free αCSF1R conditions, though not significantly different from free αPD-L1 or either single targeted liposome (all AB/A).

#### U87MG-TCM TAM2Ms

Overall division percentages were lower than those observed under A172-TCM conditions. Untreated and PEG-Lipo groups showed the lowest proliferation (D and CD). Free antibodies produced intermediate levels of T-cell restoration with αPD-L1 performing somewhat better than αCSF1R (BC for αPD-L1; C for αCSF1R). αPD-L1-Lipo and the dual combination formed the top tier of responses (AB and A, respectively); the combination was significantly higher than untreated and PEG-Lipo groups, but not significantly different from αPD-L1-Lipo alone.

#### DAOY-TCM TAM2Ms

T-cell proliferation was generally modest across all treatment conditions. Most single-agent conditions, whether free antibody or liposomal formulations, were grouped as AB and were not significantly different from PEG-Lipo. Only the dual αPD-L1+αCSF1R-Lipo combination reached the highest response group (A) and was significantly greater than untreated control (B), though not significantly different from single-agent groups (AB).

#### ONS-76-TCM TAM2Ms

Untreated and PEG-Lipo groups showed the lowest proliferation (D and D). αCSF1R and αCSF1R-Lipo produced intermediate responses (CD and BC), while free αPD-L1 performed somewhat better (BC). The highest responses were achieved with αPD-L1-Lipo (AB) and the dual combination (A); the combination was significantly greater than untreated, PEG-Lipo, and αCSF1R-based treatments, but not significantly different from αPD-L1-Lipo alone. Taken together, these data demonstrate that targeted immunoliposomes improve T-cell proliferative rescue compared with non-targeted controls, with the αPD-L1-Lipo + αCSF1R-Lipo combination consistently yielding the highest or near-highest response across all TCM conditions. The magnitude of proliferative rescue varied by TCM source, and in several contexts, the combination performed comparably to αPD-L1-Lipo rather than demonstrating strict superiority. Representative CFSE histograms are provided in Figures S4–S5.

### Liposomal Co-Targeting of PD-L1 and CSF1R on TAM2Ms Reduces Early Apoptosis of Co-Cultured Jurkat T Cells

Activated Jurkat T-cells were co-cultured for 72 h with TAM2Ms generated using TCM from A172, U87MG, DAOY or ONS-76. Early apoptosis was quantified as the Annexin V⁺/PI⁻ fraction by flow cytometry across seven conditions: untreated, PEG-Lipo, free αPD-L1, free αCSF1R, αPD-L1-Lipo, αCSF1R-Lipo, and αPD-L1+αCSF1R-Lipo (Fig. 5; representative dot plots in Fig. S6–S7).

#### A172-TCM

Untreated co-cultures exhibited the highest proportion of early apoptotic T cells. PEG-Lipo produced only a modest reduction. Free αPD-L1 reduced apoptosis to a greater extent than free αCSF1R (D vs C). The lowest apoptosis levels were observed with αPD-L1-Lipo and the dual αPD-L1-Lipo + αCSF1R-Lipo combination (E), both of which were significantly lower than untreated, PEG-Lipo, and free antibody conditions; αCSF1R-Lipo showed an intermediate effect (D–E).

#### U87MG-TCM

Baseline apoptosis was relatively low. PEG-Lipo had no significant effect (A). produced a meaningful reduction in apoptosis (D–E), whereas free αCSF1R showed only a modest effect (B–C). αPD-L1-Lipo achieved the lowest apoptosis levels of all conditions (E). The dual combination performed comparably to free αPD-L1 (D–E) but suppressed αPD-L1-Lipo. αCSF1R-Lipo outperformed its free antibody counterpart (C– D).

#### DAOY-TCM

Untreated co-cultures showed high levels of T-cell apoptosis. PEG-Lipo had a limited protective effect (B). Free αPD-L1 reduced apoptosis (C), while free αCSF1R did not produce a significant change (group B). The greatest reduction was achieved with αPD-L1-Lipo (E), with αCSF1R-Lipo showing an intermediate effect (C–D). The dual combination (D–E) reduced apoptosis relative to controls but did not outperform αPD-L1-Lipo alone.

#### ONS-76-TCM

Overall apoptosis levels were the lowest across all four models. Untreated, PEG-Lipo and both free antibodies clustered together with no significant differences among them (A). All three targeted liposomal formulations—αPD-L1-Lipo, αCSF1R-Lipo and the dual combination—significantly reduced early apoptosis to a comparable extent (B), with no significant differences observed between them.

Across TCM conditions, targeted immunoliposomes consistently reduced early T-cell apoptosis relative to untreated and PEG-Lipo controls. αPD-L1-Lipo provided the most consistent single-agent protection across models, while αCSF1R-Lipo outperformed its free antibody counterpart in all conditions. The dual αPD-L1-Lipo +αCSF1R-Lipo combination frequently ranked among the best-performing conditions but was not uniformly superior to αPD-L1-Lipo alone, suggesting context-dependent contributions of CSF1R targeting to T-cell survival.

### Liposomal Co-Targeting of PD-L1 and CSF1R on TAM2Ms Enhances Jurkat T-cell Migration Toward Brain-Tumor Spheroids

Activated, CFSE-labeled Jurkat T-cells were co-cultured with pretreated TAM2Ms and subsequently placed in the upper chamber of Cultrex-coated Transwell inserts above corresponding tumor spheroids. Migrated CFSE⁺ T cells toward the lower chamber were quantified across seven treatment conditions (Fig. 4; representative micrographs in Figures S8–S9). Groups sharing the same letter did not differ significantly at p < 0.05.

**Figure 4.**
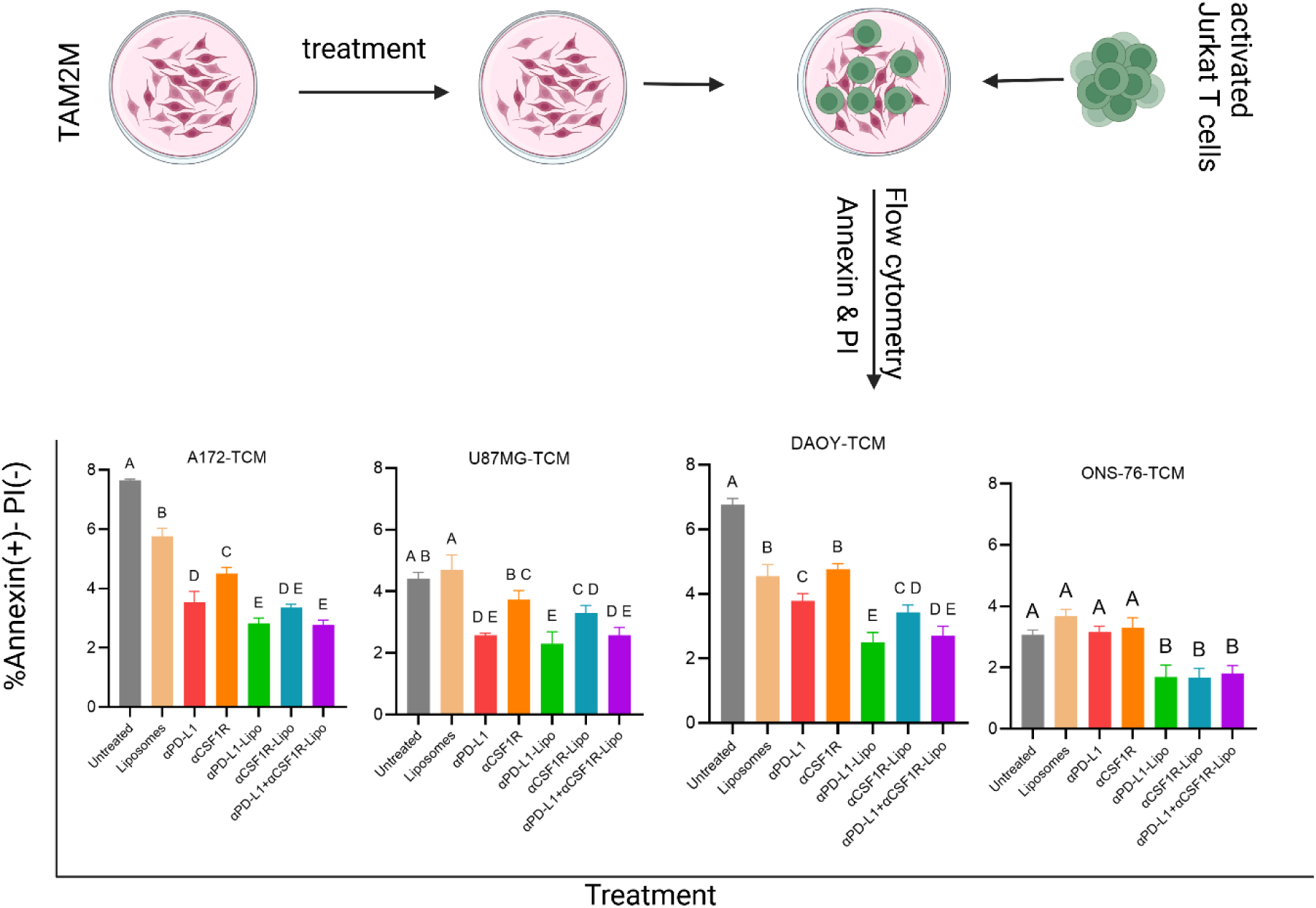
Liposomal co-targeting of PD-L1 and CSF1R on TAM2Ms reduces early apoptosis in co-cultured Jurkat T-cells across GBM- and MB-conditioned medium models. (Top) Schematic representation of the experimental workflow. T-cell apoptosis was subsequently assessed by Annexin V-FITC/PI dual staining via flow cytometry. (Bottom) Bar graphs depict the percentage of early apoptotic Jurkat T-cells (Annexin V⁺/PI⁻) with pretreated TAM2Ms under four TCM conditions: A172-TCM (GBM), U87MG-TCM (GBM), DAOY-TCM (MB), and ONS-76-TCM (MB). Treatment groups include Untreated, PEG-Lipo, free αPD-L1 nanobody (αPD-L1), free αCSF1R antibody (αCSF1R), αPD-L1-Lipo, αCSF1R-Lipo, and dual combination αPD-L1-Lipo+αCSF1R-Lipo. Data are expressed as mean ± SEM (n = 3). Different letters above bars indicate statistically significant differences between groups, as determined by one-way ANOVA with Tukey’s post hoc test (p < 0.05). Groups sharing the same letter are not significantly different from one another.

**Figure 5.**
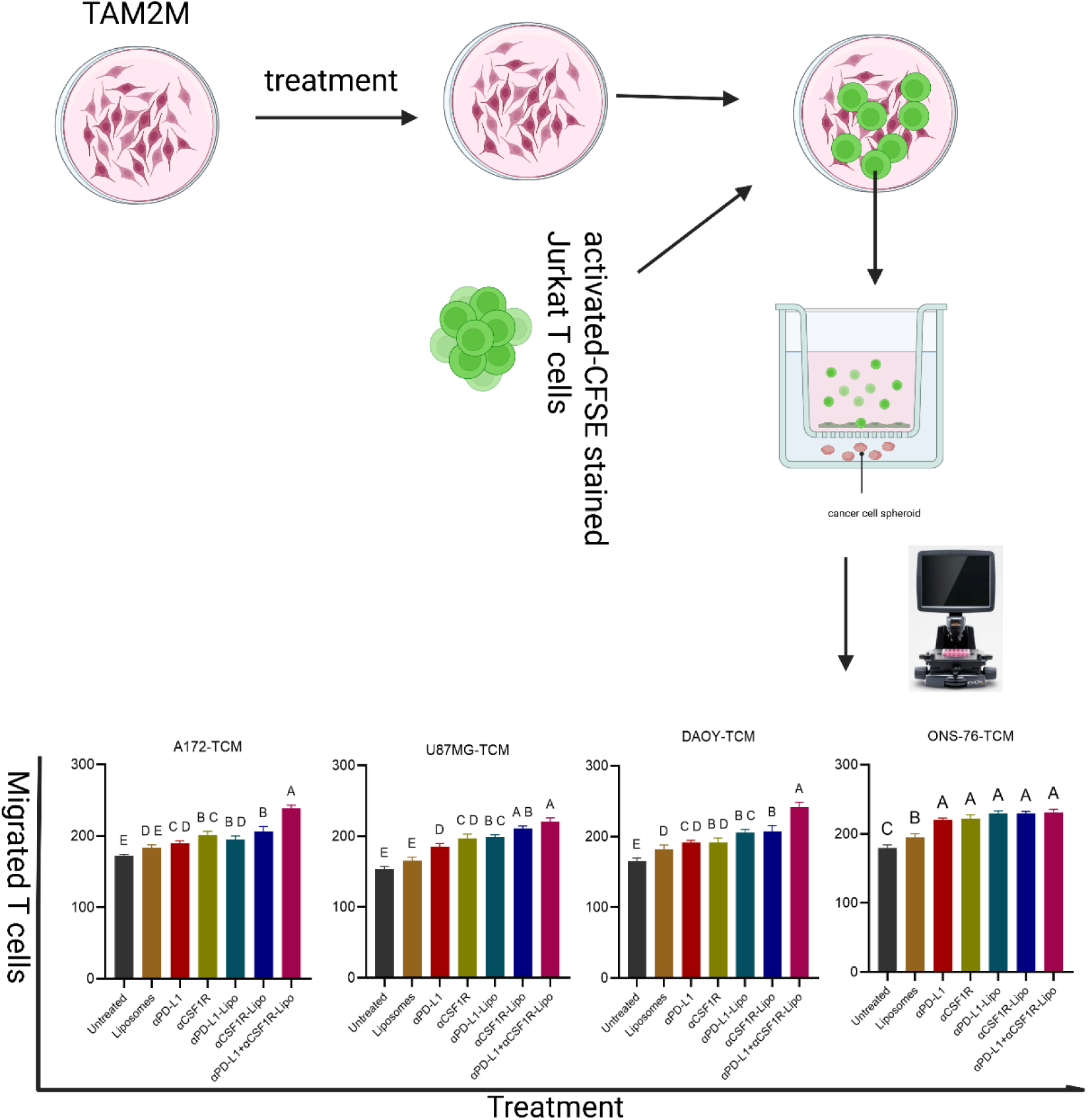
Liposomal co-targeting of PD-L1 and CSF1R on TAM2Ms enhances Jurkat T-cell migration toward brain tumor spheroids. (Top) Schematic representation of the experimental workflow. (Bottom) Bar graphs depicting the number of CFSE⁺ T cells that migrated through the Cultrex matrix toward tumor spheroids under four TCM conditions: A172-TCM (GBM), U87MG-TCM (GBM), DAOY-TCM (MB), and ONS-76-TCM (MB). Treatment groups include Untreated, PEG-Lipo, free αPD-L1 nanobody (αPD-L1), free αCSF1R antibody (αCSF1R), αPD-L1-Lipo, αCSF1R-Lipo, and dual combination αPD-L1-Lipo+αCSF1R-Lipo. Data are expressed as mean ± SEM (n= 3). Different letters above bars indicate statistically significant differences between groups as determined by one-way ANOVA with Tukey’s post hoc test (p < 0.05). Groups sharing the same letter are not significantly different from one another.

#### A172-TCM

Untreated co-cultures yielded the lowest T-cell migration. PEG-Lipo produced only a marginal increase and remained among the lower-performing groups (D–E). Free αPD-L1 and free αCSF1R provided modest improvements in migration (C and B–C). Targeted liposomal formulations (αPD-L1-Lipo and αCSF1R-Lipo) performed better than or comparably to their respective free antibody counterparts (B–D). The dual αPD-L1-Lipo+αCSF1R-Lipo combination achieved the highest migration and was significantly greater than that of all other conditions (A).

#### U87MG-TCM

A similar treatment hierarchy was observed, with generally lower absolute migration counts compared to A172-TCM. Untreated and PEG-Lipo groups showed the lowest migration (E). Free αPD-L1 and free αCSF1R produced intermediate improvements (D and C–D), while αPD-L1-Lipo and αCSF1R-Lipo further increased migration (B–C and A–B). The dual combination again produced the highest response (A), significantly exceeding untreated and PEG-Lipo conditions.

#### DAOY-TCM

Untreated co-cultures showed the lowest migration (D); PEG-Lipo showed a small but significant improvement (C). Free αPD-L1 and free αCSF1R provided additional gains (C–D and B–C), and both single-targeted liposomal formulations further increased migration (B). The dual αPD-L1-Lipo + αCSF1R-Lipo combination produced the greatest migration among all conditions and was significantly higher than all other treatment groups (A).

#### ONS-76-TCM

Baseline migration was higher in this model compared to the other TCM conditions. Untreated co-cultures showed the lowest migration (C), with PEG-Lipo performing intermediately (B). All antibody-containing treatments—free αPD-L1, free αCSF1R, αPD-L1-Lipo, αCSF1R-Lipo, and the dual combination—clustered together at the top (A) with no significant differences among them.

## Discussion and Conclusion

### Discussion

We describe a modular liposomal platform decorated with an αPD-L1 nanobody and an αCSF1R antibody (Cabiralizumab) to counteract macrophage-driven immunosuppression and restore T-cell function in brain-tumor–conditioned environments. Liposomal nanoparticles are a well-established platform with multiple clinically approved formulations, and PEGylated immunoliposomes have been widely explored for targeted delivery to tumor-associated immune cells [62]. The nanocarriers were well characterized physiochemically: all formulations fell within the 120–145 nm size range with moderately negative zeta potentials (about −30 to −40 mV), a profile associated with favorable colloidal stability and reduced nonspecific protein adsorption [63]. TEM confirmed the expected flattened vesicular morphology of phospholipid bilayers, and SDS-PAGE with reducing conditions verified successful surface conjugation of both targeting ligands. Together with the receptor-dependent uptake observed in TAM2Ms, these data suggest that the functional effects seen in co-culture reflect on-target engagement rather than nonspecific uptake or cytotoxicity. Dose–response testing further established a broad safety window, with the working dose used in functional assays remaining well below IC50 values across all TCM conditions.

Across four independent tumor-conditioned media (TCM), THP-1–derived macrophages upregulated PD-L1 and CD163, consistent with polarization toward an M2-like, immunosuppressive phenotype, in line with previously reported responses to tumor-secreted factors, including IL-4, IL-13, and CSF-1 (63). CSF1R induction was robust under U87MG, DAOY and ONS-76 TCM conditions but showed only a non-significant trend under A172-TCM, suggesting heterogeneity in CSF1R-axis engagement among tumor secretum’s. This variability likely explains why PD-L1–directed strategies were consistently effective, whereas the additional benefit of CSF1R blockade appeared to depend on the degree of CSF1R upregulation in the specific TCM context.

Functionally, targeted immunoliposomes improved three core T-cell readouts— proliferation, survival, and chemotaxis toward tumor spheroids. αPD-L1-Lipo was the most consistently effective single-agent formulation across all models, and in more immunosuppressive settings (A172-TCM and DAOY-TCM), the dual αPD-L1-Lipo + αCSF1R-Lipo combination delivered the strongest proliferative rescue. Reductions in early T-cell apoptosis paralleled these trends, with αPD-L1-Lipo providing reliable protection across conditions and αCSF1R-Lipo outperforming free αCSF1R antibody, particularly in the DAOY model. The superiority of the CSF1R-targeted liposome over its free-antibody counterpart is consistent with multivalent presentation and avidity-driven receptor engagement, which have been reported to enhance functional antagonism of surface receptors on immune cells [64]. Restoration of T-cell chemotaxis toward tumor spheroids reinforced these findings. In three of four TCM models (A172, U87MG, and DAOY), antibody-functionalized liposomes enhanced migration, and the dual combination frequently ranked highest. The ONS-76 model was an exception, as baseline migration was relatively high, and all antibody conditions—free or liposomal— performed comparably, suggesting either weaker TAM2M-mediated migratory suppression in this context or a ceiling effect of the assay. Collectively, these results highlight the context-dependent nature of dual targeting, with the greatest benefits observed when both PD-L1 and CSF1R pathways are substantively engaged by the tumor secretum.

Mechanistically, PD-L1 blockade on TAMs likely rescues T-cell receptor signaling and IL-2–driven proliferation, directly improving T-cell survival and division, consistent with established mechanisms of PD-1/PD-L1 checkpoint inhibition [65]. CSF1R inhibition may act more indirectly by reducing M2 polarization and the downstream production of immunosuppressive cytokines and chemokines that restrain T-cell function [43] . The receptor-mediated internalization of both immunoliposomes by TAM2Ms is consistent with ligand-induced clustering and down-modulation, offering a plausible mechanistic basis for the enhanced activity of liposomal relative to free CSF1R antibody [66]Differences in treatment responses between GBM and MB models—and between A172 and U87MG within GBM—likely reflect distinct secretum compositions that differentially modulate these immunosuppressive axes.

Several limitations of this study warrant consideration. THP-1–derived macrophages and Jurkat T-cells provide reproducible and well-controlled model systems, but do not fully recapitulate the phenotypic diversity, exhaustion states, or functional heterogeneity of primary human tumor-infiltrating immune cells [67,68] . The migration assay, while incorporating three-dimensional cancer cell spheroids and basement membrane extract to better approximate the tumor microenvironment, cannot replicate the complexity of in vivo stromal and vascular barriers. Statistical groupings demonstrate significance between conditions but do not establish causality or define the molecular pathways underlying the observed effects. Furthermore, the combination condition in this study involved the co-administration of two separately targeted liposomal formulations; whether a single liposome decorated with both antibodies would further improve efficacy and alter receptor engagement dynamics remains to be investigated.

These findings support the concept of TAM-targeted, checkpoint-blocking nano therapy as a strategy to normalize immune function in brain-tumor microenvironments. The consistent potency of αPD-L1-Lipo, the context-dependent contribution of CSF1R blockade, and the frequent superiority of the dual combination in more immunosuppressive settings collectively suggest that the therapeutic benefit of this approach may be greatest in tumors with high TAM2M infiltration and active engagement of both the PD-L1 and CSF1R axes. Future studies should investigate the cytokine and chemokine programs induced in TCM-educated TAMs following treatment, assess macrophage repolarization at the phenotypic and transcriptional levels, and validate key findings in primary human macrophage and T-cell systems. Additional work should compare dual-decorated single-nanoparticle formulations with antibody mixture formats and optimize ligand density and antibody ratios to maximize receptor co-engagement. Finally, evaluation of pharmacokinetics, tumor accumulation, immune cell infiltration, and therapeutic efficacy in orthotopic GBM and MB animal models will be essential steps toward translational development of this platform.

## Conclusion

We engineered stable PEGylated liposomes bearing an αPD-L1 nanobody and an αCSF1R antibody that selectively engage and are internalized by tumor-conditioned, M2-like macrophages. Brain tumor TCMs derived from GBM and MB cell lines upregulated PD-L1, CSF1R, and CD163 on THP-1-derived macrophages, confirming the availability of both therapeutic targets under tumor-relevant conditions. At non-toxic doses, the targeted immunoliposomes restored multiple T-cell functions in co-culture, enhancing proliferation, reducing early apoptosis, and improving migration toward tumor spheroids. αPD-L1-Lipo was the most consistently effective single-agent formulation across all models, while the dual αPD-L1-Lipo + αCSF1R-Lipo combination frequently yielded the strongest responses, particularly in the more immunosuppressive A172-TCM and DAOY-TCM settings, suggesting context-dependent cooperativity between these two pathways. Taken together, these results support macrophage-directed, checkpoint-blocking nano therapy as a potential strategy to normalize the immunosuppressive milieu in brain tumors and provide a rationale for further optimization and *in vivo* validation, including evaluation of dual-decorated single-carrier formulations, systematic tuning of ligand density and antibody ratios, and stratification of treatment approaches according to TAM phenotype and tumor secretum composition.

## Supporting information

Additional supporting information is available in the supporting information section.

## Author contributions

Ommolbanin Asad Pour contributed to conceptualization, funding acquisition, investigation, formal analysis, visualization, and writing (original draft and editing). Aylar Asadpour contributed to preparing the graphical image. Jamal Ghanam and Venkatesh Kumar Chetty contributed to methodology. Fatemeh Rahbarizadeh contributed to liposome preparation, perspective, interpretation of liposome characterization, and critical revision of the manuscript. Susann Hetze, Lennart Barthel, and Zohreh Amozgar contributed to the translational neuro-oncology perspective, interpretation of glioblastoma-relevant aspects, and critical revision of the manuscript. Basant Kumar Thakur contributed to supervision (project oversight), and all the authors contributed to writing (review and editing).

## Data availability

The data presented in this study are available within the article and its supplementary information files.

## Declarations

### Ethics approval and consent to participate

Not applicable

### Consent for publication

The authors declare no competing financial interests or personal relationships that could have influenced the work reported in this paper.

### Use of generative AI and AI-assisted technologies

Perplexity AI and Grammarly were used solely for grammar and language editing of the final manuscript draft. All authors reviewed and verified the accuracy, originality, and scientific content. No AI tool contributed to study design, data generation, analysis, interpretation, or creation of figures/tables.

## Acknowledgements

All authors thank Prof. Felipe Opazo for providing the anti-PD-L1 nanobody, and Prof. Bernd Giebel, Institute for Transfusion Medicine, University Hospital Essen, for providing access to the Ultracentrifugation and NTA device in his laboratory. We thank the Imaging Center Essen (IMCES) at the Faculty of Medicine of the University of Duisburg-Essen, Germany, for providing access to its electron microscopy services and especially Sylvia Voortmann for sample preparation and Bernd Walkenfort for transmission and scanning electron microscopy. We also thank IMCES for providing access to the Zeiss Elyra PS.1

