## Supplementary material for "Dual Immunoliposome Targeting of PD-L1 and CSF1R affects T-Cell readouts in tumor-conditioned co-cultures: An *In Vitro* Study in Glioblastoma and Medulloblastoma": Supplemantry figures

### Supplementary Figures

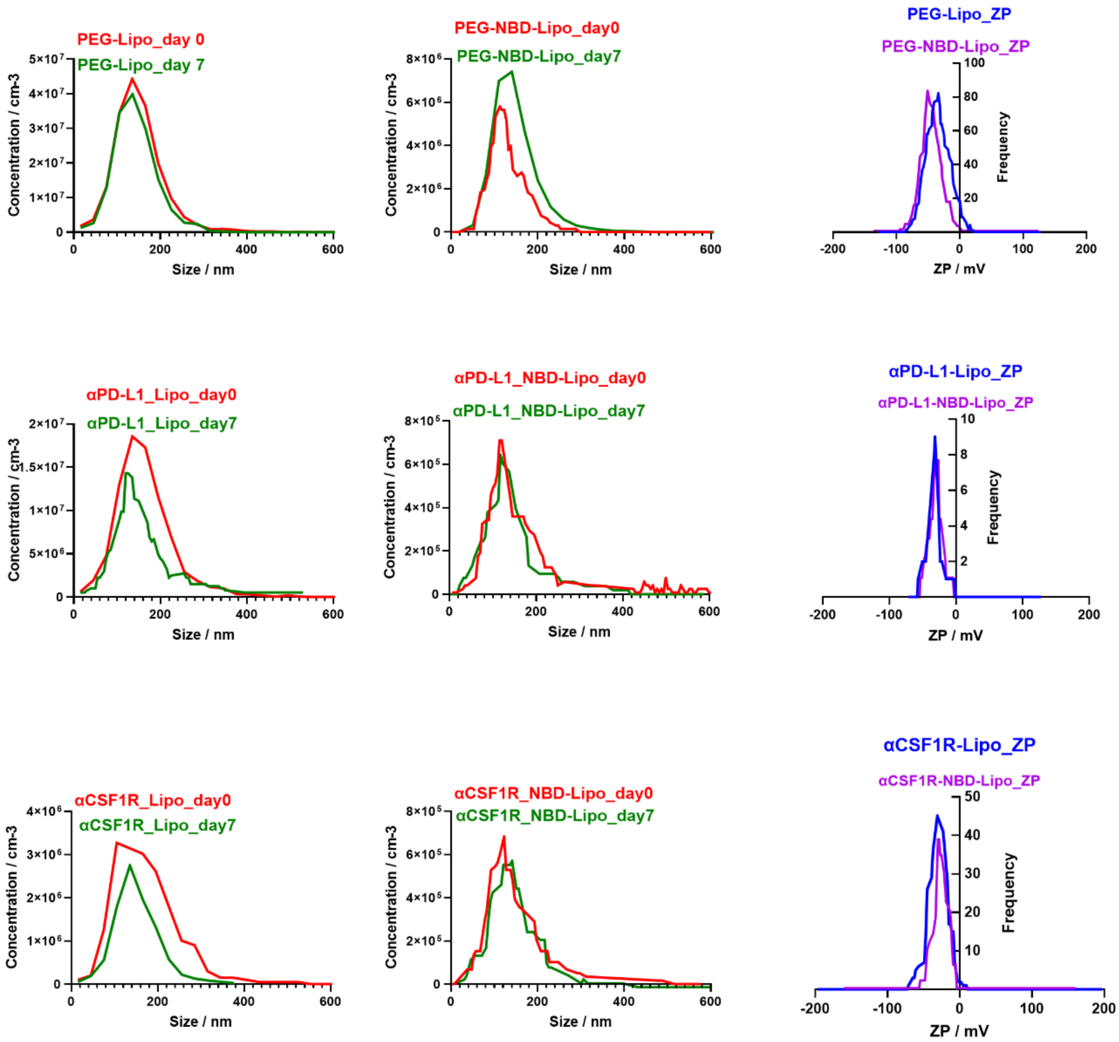


**Figure S1. NTA size distribution profiles and zeta potential measurements of liposomal formulations**. Nanoparticle tracking analysis (NTA) size distribution profiles (left and center columns) of PEG-Lipo, PEG-NBD-Lipo (Row 1), αPD-L1-Lipo, αPD-L1-NBD-Lipo (Row 2), αCSF1R-Lipo, and αCSF1R-NBD-Lipo (Row 3) measured at day 0 (red) and day 7 (green) of storage at 4°C. Profiles plotted as particle concentration (particles/cm$^3$) versus hydrodynamic diameter (nm). Zeta potential (ZP) frequency distributions (right column) of non-fluorescent (blue) and NBD-labeled (magenta) variants of each formulation at day 0, plotted as frequency versus zeta potential (mV).

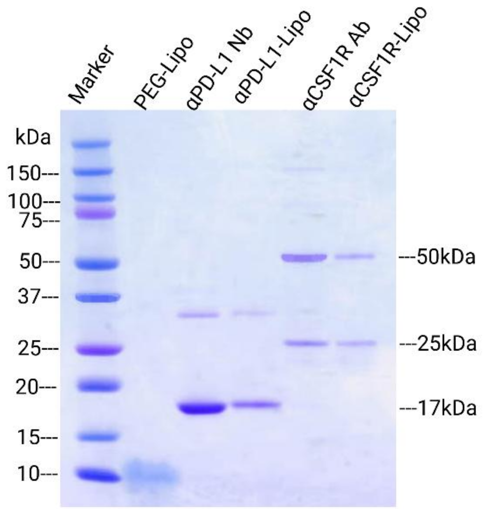


**Figure S2. SDS-PAGE analysis confirming the surface conjugation of the anti-PD-L1 nanobody and the anti-CSF1R antibody to liposomal nanoparticles.** Reducing SDS-PAGE of PEG-Lipo (non-functionalized liposome control), free anti-PD-L1 nanobody (αPD-L1 Nb), αPD-L1-Lipo, free anti-CSF1R antibody (αCSF1R Ab; Cabiralizumab), and αCSF1R-Lipo. A prestained molecular weight marker (10–150 kDa) was included for molecular weight reference. Proteins were visualized by Coomassie Brilliant Blue R-250 staining. Key bands: αPD-L1 Nb monomer (~17 kDa); αCSF1R Ab heavy chain (~50 kDa); αCSF1R Ab light chain (~25 kDa).

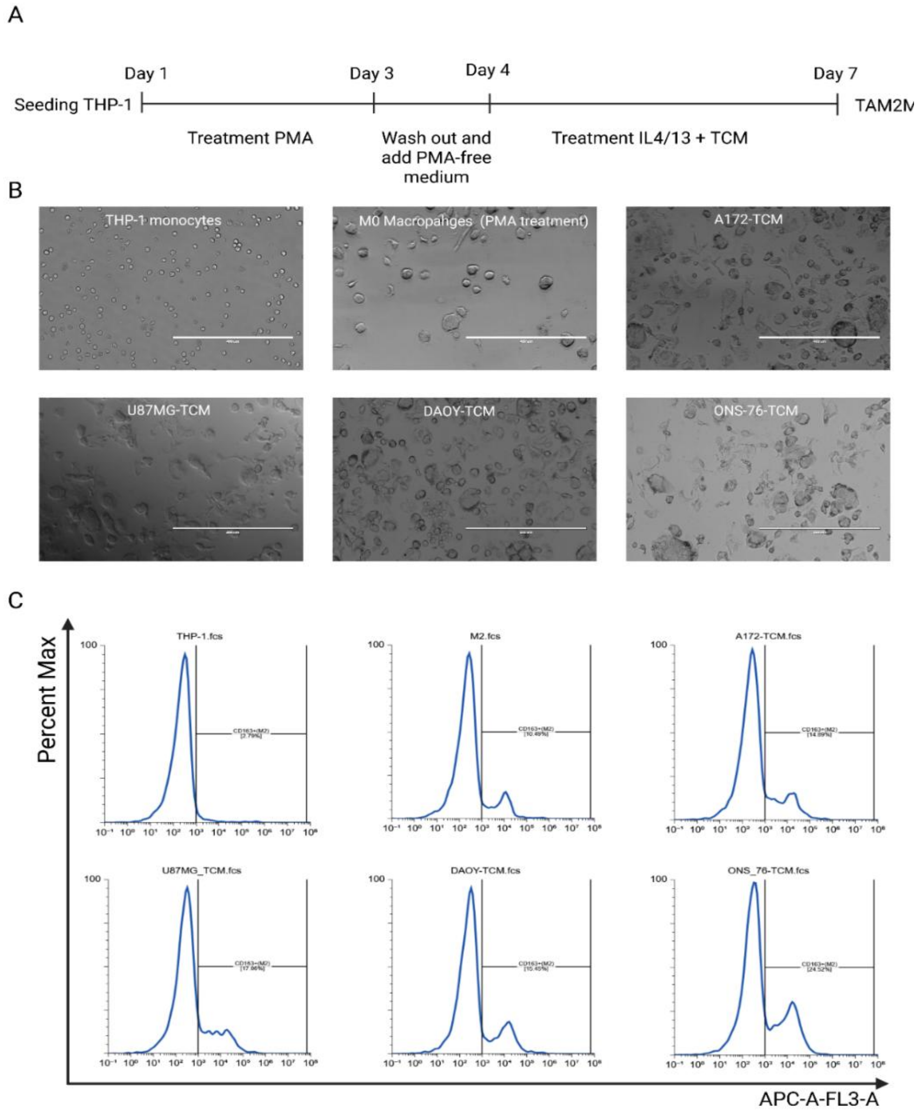


**Figure S3. THP-1 monocyte differentiation and polarization into TAM2Ms: timeline, morphology, and CD163 expression.** (A) Schematic timeline of the TAM2M generation protocol, illustrating sequential stages of PMA-induced differentiation and subsequent TCM/cytokine-driven M2 polarization. (B) Representative phase-contrast micrographs of THP-1 monocytes, PMA-differentiated M0 macrophages, and TAM2Ms polarized under A172-TCM, U87MG-TCM, DAOY-TCM, and ONS-76-TCM conditions. Scale bars as indicated are 400 μm; Images acquired at 10× magnification (EVOS FL). (C) Representative flow cytometry histograms of CD163 surface expression (APC channel) across undifferentiated THP-1 monocytes, IL-4/IL-13-polarized macrophages (M2-IL4/13), and

TCM-conditioned TAM2Ms generated under each tumor cell line condition–x-axis: APC fluorescence intensity; y-axis: percent of maximum.

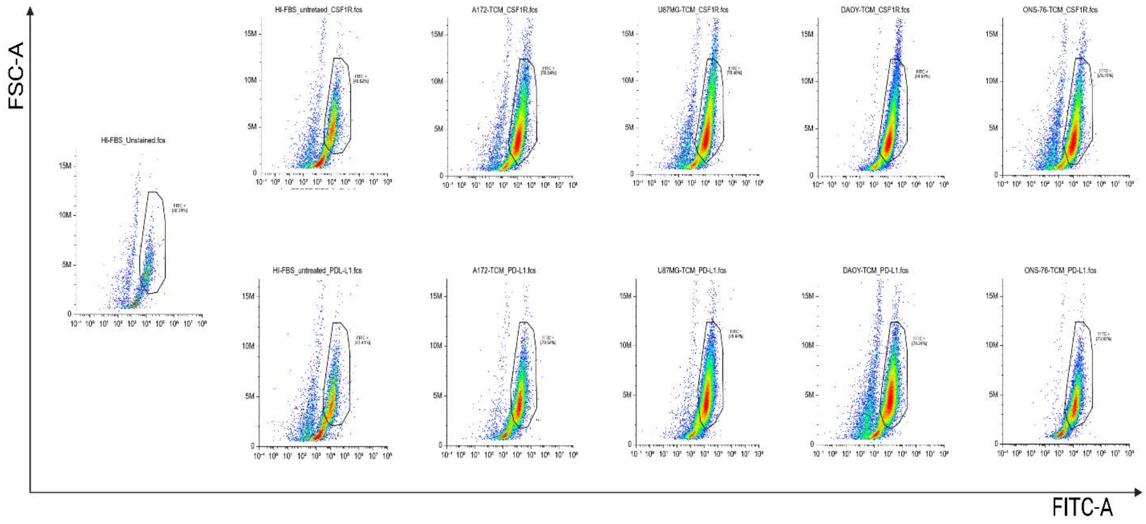


**Figure S4. Flow cytometric analysis of CSF1R and PD-L1 surface expression on TAM2Ms.** Representative FSC-A versus FITC-A dot plots showing CSF1R (top row) and PD-L1 (bottom row) surface expression on unpolarized macrophages (M0; HI-FBS untreated) and TAM2Ms polarized under A172-TCM, U87MG-TCM, DAOY-TCM, and ONS-76-TCM conditions. An unstained control (far left) was included to establish the autofluorescence baseline. Polygon gates identify FITC-positive cell populations, with the percentage of FITC$^+$ cells indicated within each panel.

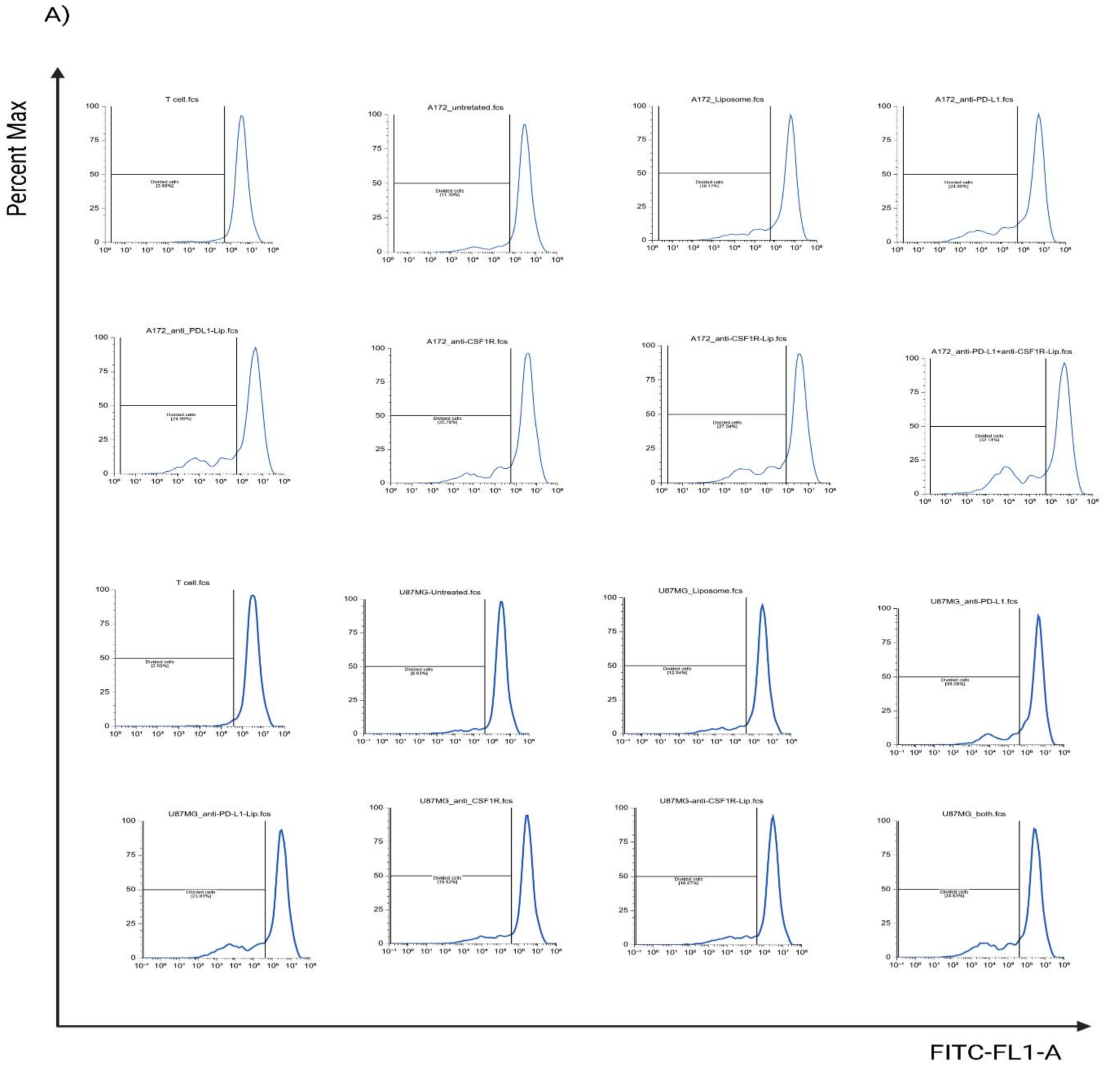


**Figure S5. Flow cytometric analysis of glioblastoma TAM2M liposomal treatments on T-cell proliferation in co-cultures.** Representative flow cytometry histograms depicting CFSE dilution as a measure of T-cell proliferation. CFSE-labelled T cells were co-cultured with TAM2M differentiated and polarized under A172-TCM (top two rows) or U87MG-TCM (bottom two rows) conditions and subjected to the following treatment groups: untreated, PEG-Lipo, free αPD-L1 Nb, αPDL1-Lipo, free αCSF1R Ab, αCSF1R-Lipo, or dual combination of αPD-L1-Lipo + αCSF1R-Lipo. The x-axis represents CFSE fluorescence intensity (FITC-FL1-A, log scale) and the y-axis represents normalized cell frequency (percent of maximum). The horizontal gate denotes the divided T-cell population, with the corresponding percentage of divided cells indicated within each histogram. Progressive leftward shifts in CFSE fluorescence intensity are indicative of increased T-cell proliferation.

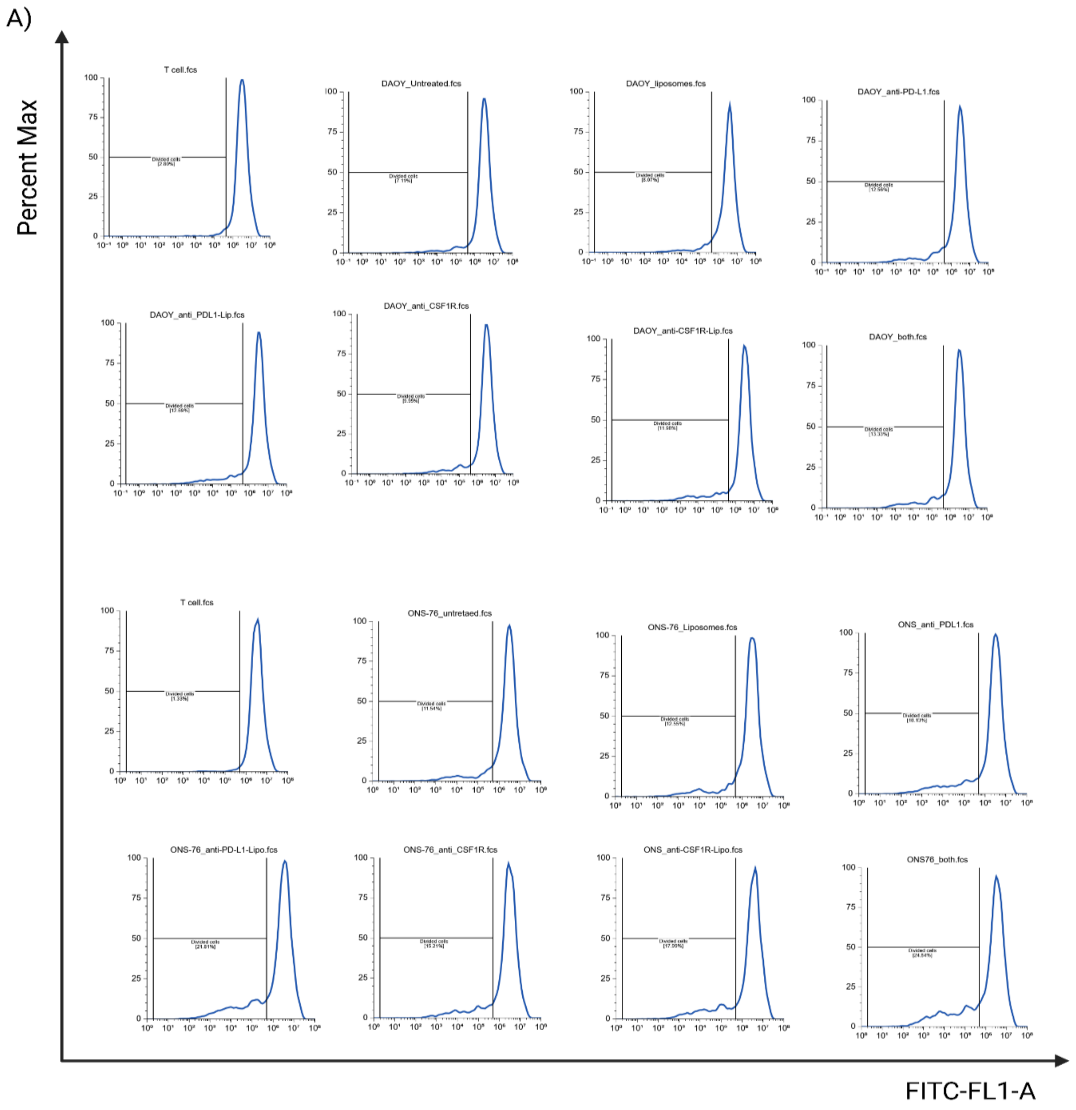


**Figure S6. Flow cytometric analysis of medulloblastoma TAM2M liposomal treatments on T-cell proliferation in co-cultures.** Representative flow cytometry histograms depicting CFSE dilution as a measure of T-cell proliferation. CFSE-labelled T cells were co-cultured with TAM2M differentiated and polarized under DAOY-TCM (top two rows) or ONS-76-TCM (bottom two rows) conditions and subjected to the following treatment groups: untreated, PEG-Lipo, free αPD-L1 Nb, αPDL1-Lipo, free αCSF1R Ab, αCSF1R-Lipo, or combination of αPD-L1-Lipo + αCSF1R-Lipo. The x-axis represents CFSE fluorescence intensity (FITC-FL1-A, log scale) and the y-axis represents normalized cell frequency (percent of maximum). The horizontal gate denotes the divided T-cell population, with the corresponding percentage of divided cells indicated within each histogram. Progressive leftward shifts in CFSE fluorescence intensity are indicative of increased T-cell proliferation.

A)

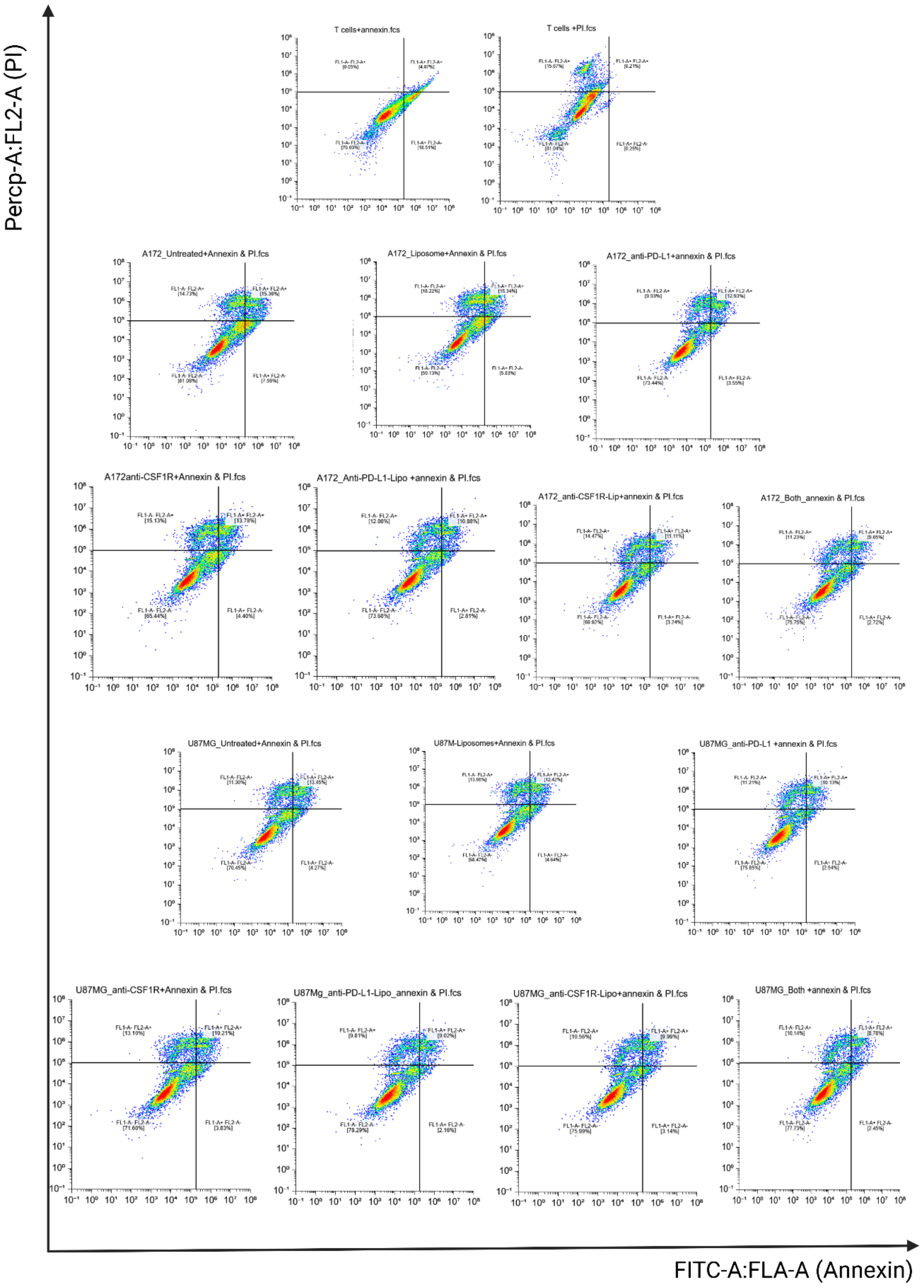
Percp-A:FL2-A (PI)
FITC-A:FLA-A (Annexin)
T cells+annexin.fcs
T cells +PI.fcs
A172_Untreated+Annexin & PI.fcs
A172_Liposome+Annexin & PI.fcs
A172_anti-PD-L1+annexin & PI.fcs
A172anti-CSF1R+Annexin & PI.fcs
A172_Anti-PD-L1-Lipo +annexin & PI.fcs
A172_anti-CSF1R-Lip+annexin & PI.fcs
A172_Both_annexin & PI.fcs
U87MG_Untreated+Annexin & PI.fcs
U87M-Liposomes+Annexin & PI.fcs
U87MG_anti-PD-L1 +annexin & PI.fcs
U87MG_anti-CSF1R+Annexin & PI.fcs
U87Mg_anti-PD-L1-Lipo_annexin & PI.fcs
U87MG_anti-CSF1R-Lipo+annexin & PI.fcs
U87MG_Both +annexin & PI.fcs

**Figure S7. Flow cytometric analysis of glioblastoma TAM2M liposomal treatments on T-cell apoptosis in co-cultures.** Representative Annexin V-FITC/PI dual-staining dot plots depicting T-cell apoptosis. T cells were co-cultured with TAM2Ms differentiated and polarized under A172-TCM (top two rows) or U87MG-TCM (bottom two rows) conditions and subjected to the following treatment groups: untreated, PEG-Lipo, free αPD-L1 nanobody (αPD-L1 Nb), αPD-L1-Lipo, free αCSF1R antibody (αCSF1R Ab), αCSF1R-Lipo, or dual combination αPD-L1-Lipo + αCSF1R-Lipo. Quadrant designations are as follows: lower left ($FL1\text{-}A^{-}/FL2\text{-}A^{-}$) = viable cells; lower right ($FL1\text{-}A^{+}/FL2\text{-}A^{-}$) = early apoptotic cells; upper right ($FL1\text{-}A^{+}/FL2\text{-}A^{+}$) = late apoptotic/secondary necrotic cells; upper left ($FL1\text{-}A^{-}/FL2\text{-}A^{+}$) = necrotic cells. The percentage of cells within each quadrant is indicated. Data are representative of at least three independent experiments.

A)

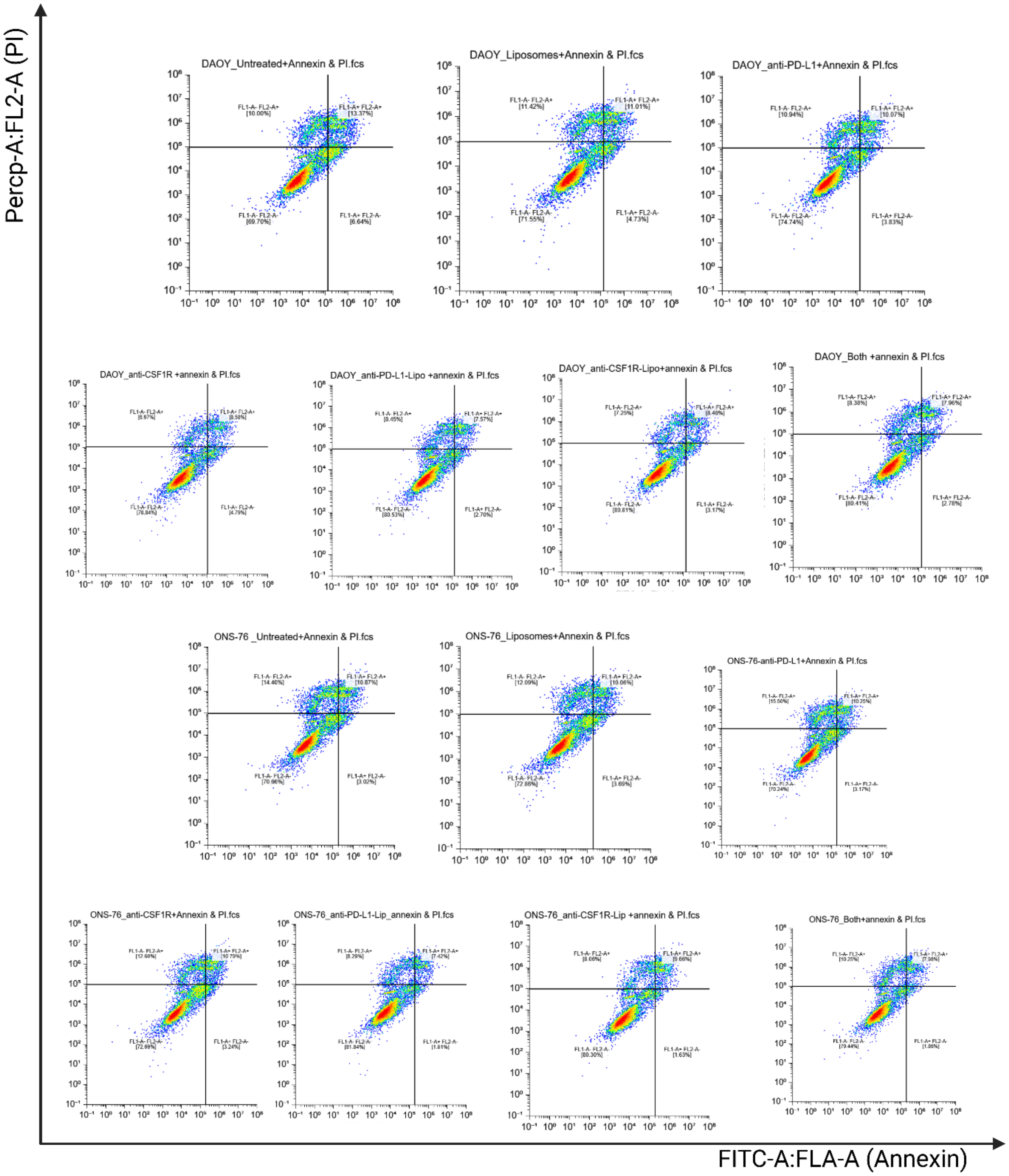


**Figure S8. Flow cytometric analysis of medulloblastoma TAM2M liposomal treatments on T-cell apoptosis in co-cultures.** Representative Annexin V-FITC/PI dual-staining dot plots depicting T-cell apoptosis. T cells were co-cultured with TAM2Ms differentiated and polarized under DAOY-TCM (top two rows) or ONS-76-TCM (bottom two rows) conditions and subjected to the following treatment groups: untreated, PEG-Lipo, free αPD-L1 nanobody (αPD-L1 Nb), αPD-L1-Lipo, free αCSF1R antibody (αCSF1R Ab), αCSF1R-Lipo, or dual combination αPD-L1-Lipo + αCSF1R-Lipo. Quadrant designations are as follows: lower left (FL1-$A^-$/FL2-$A^-$) = viable cells; lower right (FL1-$A^+$/FL2-$A^-$) = early apoptotic cells; upper right (FL1-$A^+$/FL2-$A^+$) = late apoptotic/secondary necrotic cells; upper left (FL1-$A^-$/FL2-$A^+$) = necrotic cells. The percentage of cells within each quadrant is indicated. Data are representative of at least three independent experiments.

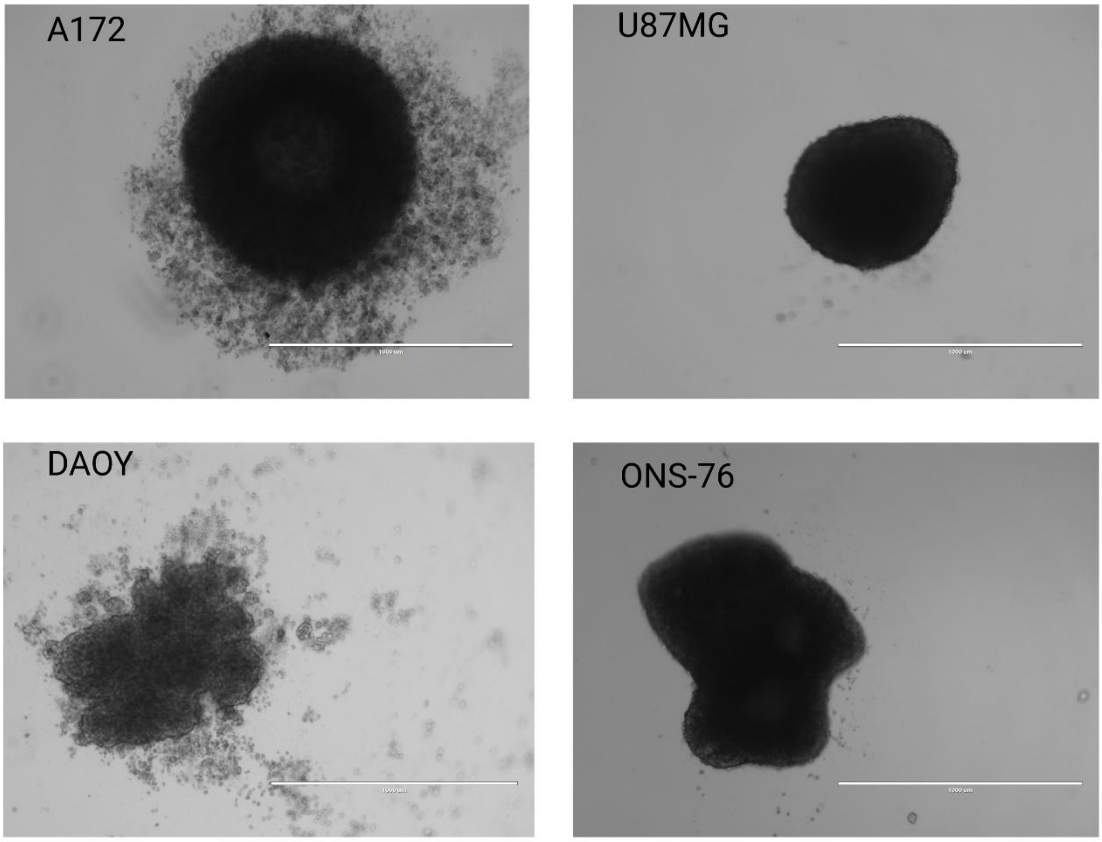


**Figure S9. Representative microscopy images of spheroids generated from glioblastoma and medulloblastoma cell lines.** Representative brightfield images of three-dimensional (3D) tumor spheroids formed from two glioblastoma cell lines (A172 and U87MG) and two medulloblastoma cell lines (DAOY and ONS-76), generated under ultra-low attachment conditions in serum-free spheroid medium. Images were acquired using inverted microscopy following 48 hours of self-assembly. Scale bars = 1000 μm.

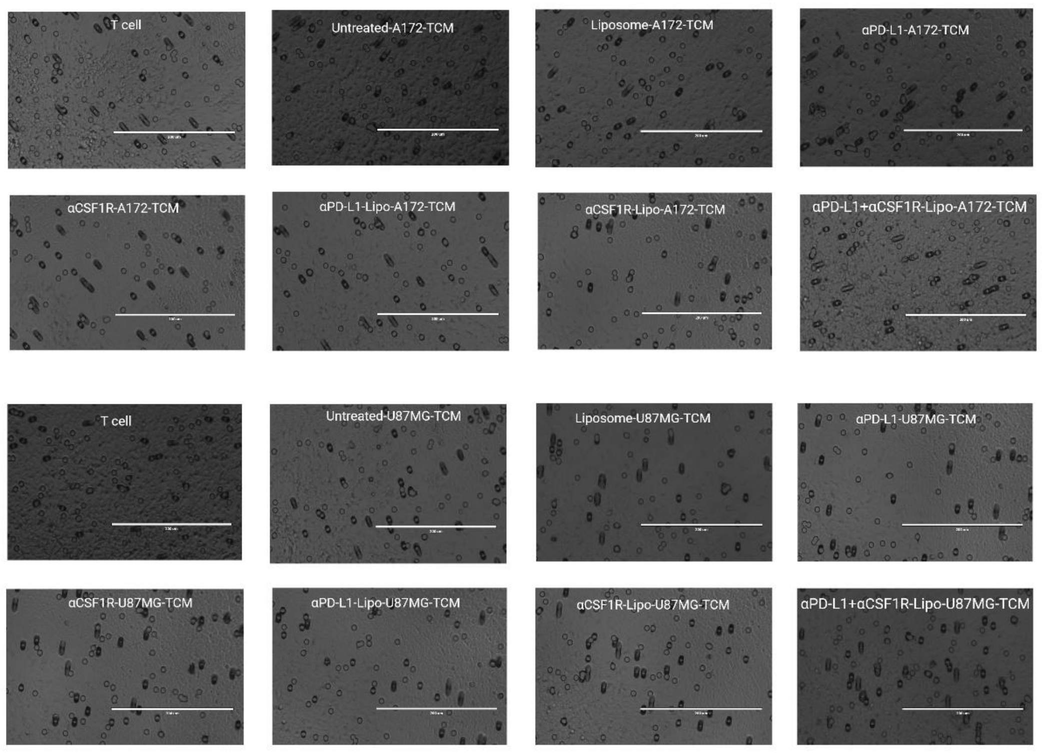


**Figure S10. Microscopy images of T-cell migration toward glioblastoma cell line spheroids.** Representative brightfield micrographs of CFSE-labeled T cells that migrated into the Cultrex basement membrane extract (BME) layer on the upper surface of Transwell membranes in response to a chemoattractant gradient established by tumor spheroids placed in the lower chamber. Images were acquired at consistent magnification across all conditions to enable comparison of T-cell invasion within the gel matrix. Scale bars = 200 μm.

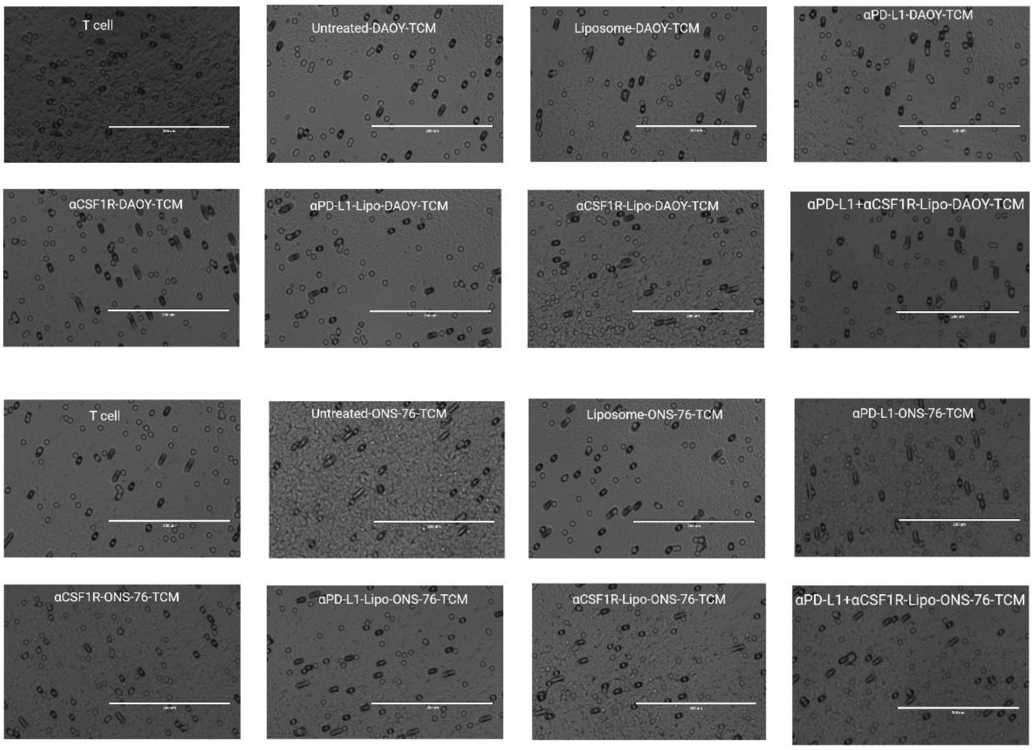


**Figure S11. Microscopy images of T-cell migration toward medulloblastoma cell line spheroids.** Representative bright-field micrographs of T cells that migrated into the Cultrex (basement membrane extract) layer on the upper surface of Transwell membranes when exposed to a cancer cell spheroid on the lower surface. Images were acquired at the same magnification to visualize cell invasion within the gel. Scale bars, 200 μm.
